# Polygenic basis of Bt Cry resistance evolution in wild *Helicoverpa zea*

**DOI:** 10.1101/2024.02.09.579652

**Authors:** Katherine L Taylor, Jane Quackenbush, Cara Lamberty, Kelly A Hamby, Megan L Fritz

## Abstract

Strong and shifting selective pressures of the anthropocene are rapidly shaping phenomes and genomes of organisms worldwide. One major selective force on insect genomes is crops expressing pesticidal proteins from *Bacillus thuringiensis* (Bt). Here we characterize a rapid response to selection by Bt crops in a major crop pest, *Helicoverpa zea*. We reveal the polygenic architecture of Bt resistance evolution in *H. zea* and identify multiple genomic regions underlying this trait. In the genomic region of largest effect, we identified a gene cluster amplification, where resistant individuals showed variation in copy number for multiple genes. Signals of this amplification increased over time, consistent with the history of field-evolved Bt resistance evolution. Modern wild populations from disparate geographical regions are positive for this variant at high, but not fixed, allele frequencies. We also detected selection against single copy variants at this locus in wild *H. zea* collected from Bt expressing plants, further supporting its role in resistance. Seven trypsin genes were present in this genomic region and all appeared to be significantly upregulated in Bt resistant *H. zea*. Biochemically inhibiting trypsin activity decreased *H. zea*’s tolerance to Bt. These findings characterize rapid genome evolution in a major crop pest following anthropogenic selection and highlight the role that gene copy number variants can have in rapid evolutionary responses.

## Introduction

The study of evolution in the Anthropocene has revealed that adaptive organismal responses can occur on timescales much shorter than previously thought possible (Carroll et al. 2007). Human activities impose strong and shifting selection pressure on communities of organisms, shaping their phenomes and genomes (Palumbi, 2001). Empirical evidence of rapid evolutionary change is mounting (*e.g.* Bergland et al., 2014; Bi et al., 2019; Campbell-Staton et al., 2017; Chaturvedi et al., 2021; Ergon et al., 2019; Mikheyev et al., 2015; Roberts Kingman et al., 2021; Rudman et al., 2022; Schiebelhut et al., 2018; Stahlke et al., 2021; Storz & Wheat, 2010), and has linked the strength and nature of selection to specific genomic variants that enhance the fitness of wild organisms. Rapid evolution can result from allele frequency changes at a single major effect locus, or at many loci across the genome (reviewed in Bay et al., 2017). Currently, a major objective of the field of evolutionary genomics is to move beyond documenting the phenomenon of rapid evolution, and instead, uncover the rules that govern how these responses occur on short timescales. This includes characterizing the types of genomic variants (*i.e.* single nucleotide polymorphisms, insertion/deletion polymorphisms, copy number variants, epigenetic modifications) that are most critical for and facilitate rapid evolutionary responses. Perhaps even more important is understanding how fitness-conferring variants came to be; whether they arose *de novo*, were introduced through migration, or were selected from standing genetic variation.

Agricultural ecosystems (agroecosystems) are useful for investigating mechanisms of rapid evolution, both due to the well-understood nature of selection in these ecosystems and their relatively simplified ecosystem structure (Chen & Schoville, 2018). One common feature of agroecosystems is the use of population suppression practices, which exert strong pressure on so-called agricultural pests to evade management through resistance evolution. High levels of resistance can evolve rapidly (Brevik et al., 2018), and in arthropods alone, resistance to over 300 pesticidal compounds in more than 600 species has been documented (Fritz, 2022; Mota-Sanchez & Wise, 2022). Rapid pesticide resistance evolution also has economic and environmental consequences, which provides incentive for its study. Resistance evolution causes billions of dollars in crop losses every year, and often results in increased environmental pesticide inputs in an attempt to prevent these losses (Gould et al., 2018; Palumbi, 2001).

We have adopted the Bt cropping system as an experimental model for the study of rapid organismal evolution. Since 1996, Bt crops have provided area-wide pest suppression, while decreasing reliance on harmful broad-spectrum pesticides (Cattaneo et al., 2006; Dively et al., 2018; Hutchison et al., 2010; Kathage & Qaim, 2012; Perry et al., 2016). Bt crops have been engineered to express genes from the bacterium, *Bacillus thuringiensis*, which encode crystalline (Cry) and vegetative insecticidal proteins (Vip). The proteins specifically target lepidopteran and coleopteran species that damage crops, some of which have been historically challenging to manage with sprayable, synthetic pesticides. They have been widely adopted for insect management around the globe, and at present, 85% of corn and 89% of cotton planted in the United States express Bt traits (USDA-ERS, 2023). Bt crops are, therefore, a major selective force in agroecosystems (Gassmann & Reisig, 2023; Tabashnik & Carrière, 2017).

To improve the durability of these crops and slow resistance evolution, management plans strongly rooted in evolutionary theory, modeling-based studies, and past empirical observations were developed (*e.g.* Alstad & Andow, 1995; Frutos et al., 1999; Gould et al., 2018; McGaughey et al., 1998). Bt crops were designed to produce a “high dose”, or 25 times the amount of pesticidal protein needed to cause mortality in susceptible target pests (US EPA-SRP, 1998). Such a high dose was predicted to favor resistance arising from a single mutation of major effect, rather than from standing genetic variation across the genome (McKenzie & Batterham, 1994). Multiple toxins were also pyramided into Bt crops to redundantly kill pests that evolved resistance to a single protein. Finally, non-expressing plants, or a “refuge” planted nearby, should produce homozygous susceptible pests that dilute resistance alleles in a local landscape (Gould, 1998; Huang et al., 2011). Under these circumstances, if resistance was recessive and resistance alleles incurred a fitness cost in the absence of these toxic proteins, the emergence and spread of resistance may be preventable (Gould, 1998). Although there have been cases of successful resistance management in some target pests of Bt crops, practical resistance has emerged in several lepidopteran and coleopteran species, including the polyphagous pest, *Helicoverpa zea* (Tabashnik & Carrière, 2017).

*H. zea*, a pest of corn and cotton, was successfully managed by Bt crops when they were commercially released in 1996. Yet within 20 years, susceptibility to Cry proteins in *H. zea* had decreased and crop damage increased (Dively et al., 2016; Dively et al., 2023; Kaur et al., 2019; Yang et al., 2019; Yang et al., 2022). Resistance evolution in *H. zea* had been predicted because Cry expressing Bt crops do not produce a “high” dose for *H. zea* (Horner et al., 2003), and refuge implementation for corn and cotton was often insufficient (Reisig, 2017; Reisig & Kurtz, 2018). Resistance may also have been facilitated by cross pollination between Bt and refuge corn, which produces kernels expressing low Cry toxin doses (Pezzini et al., 2024; Dively et al., 2020; Yang et al., 2015). The Cry resistance that emerged in wild *H. zea* has provided an opportunity to empirically examine the mechanisms underlying rapid adaptation in this human-altered system. If *H. zea*’s Cry toxin exposure was not high dose and the strength of selection allowed for survivors within their natural viability distribution, adaptive responses should have a polygenic trait architecture (McKenzie & Batterham, 1994). Our previous work supported this prediction, using two single-family QTL analyses to characterize the molecular architecture of resistance to two separate Bt crop cultivars (Taylor et al., 2021).

Cry toxicity is generally understood to begin with solubilization and activation of the Cry protoxin by proteolytic enzymes in the insect digestive system, although this activation step is not necessary for the activated toxins expressed by many Bt crops (Clark et al., 2005; Gould, 1998; Székács et al., 2010). Activated toxins interact with a series of midgut receptors, eventually causing lysis of the midgut epithelial cells and leading to growth suppression or mortality in targeted insects (Heckel, 2020; Jurat-Fuentes et al., 2021; L. Liu et al., 2021). The multi-step and complex molecular mechanism of Cry toxicity provides many molecular and physiological pathways for the evolution of resistance. In other lepidopteran species, resistance has generally been connected to the disruption of the midgut toxin binding sites or reduced expression of target genes, although there are also examples of resistance associated with altered Bt protein processing, detoxification, and changes in immune function (Heckel, 2020; Jurat-Fuentes et al., 2021; L. Liu et al., 2021). Our prior work with *H. zea* led us to reject most of the common Cry resistance mechanisms described in other lepidopteran species, and instead pointed to novel resistance-related genomic regions (Taylor et al., 2021), with one under particularly strong selection by Cry toxins (Pezzini et al. 2024).

Here, we have expanded our analysis of the rapid adaptation to Bt crops observed in wild *H. zea*, describing its underlying genetic architecture. We provide further evidence to support a polygenic trait architecture of Cry resistance in wild *H. zea*. We also provide the first evidence that a copy number variant (CNV) containing a cluster of 10 genes plays a major role in the Cry resistance phenotype we observed in *H. zea*. Our results connect the introduction and adoption of transgenic crops in the landscape, which resulted in a resistance phenotype, to genomic targets of selection, and the nature of the genomic variants that facilitated *H. zea*’s rapid evolutionary response.

## Results

### Cry resistance and general growth phenotypes

To link regions of the *H. zea* genome to Cry resistance phenotypes, we used a replicated quantitative trait locus (QTL) analysis. This well-established approach relies on crosses between individuals from phenotypically distinct inbred lines or outbred populations to produce F2 offspring, whose chromosomes represent unique genomic combinations from cross founders (Falconer, 1996; Lynch & Walsh, 1998). Correlated traits within phenotypically distinct cross- founding populations are, therefore, uncorrelated in F2 progeny due to genome-wide recombination during meiosis. We crossed field-evolved resistant grandparents to one of very few known susceptible *H. zea* populations in our replicated design. This allowed us to test for genome-wide associations between field-relevant Cry resistance phenotypes and genotypes across recombined F2 chromosomes. While multiple genomic regions were associated with Cry resistance in our previous work (Taylor et al. 2021), testing of additional replicate families increased our power to detect loci of major effect that commonly contribute to field-evolved Cry resistance in wild *H. zea*.

We initially measured Cry resistance phenotypes for a field resistant population and a long-term laboratory-reared susceptible population, as well as the F2 hybrid offspring from 10 intercross families. Resistant individuals should be able to grow while feeding on diets containing Cry toxins, though at different rates depending on the number and effect size of their Cry resistance alleles, while susceptible individuals should not. Thus, measuring weight following 7 days exposure to Cry toxin-incorporated diet served as our metric of resistance. A critical component of our experimental design included measurement of intra-family controls, where half of the offspring from each cross were assayed on identical diets lacking Cry toxin. These intra-family controls served as a way to measure and, if necessary, exclude general growth-related QTL as candidates for resistance. If a QTL were truly related to Cry resistance, we reasoned that QTL associated with growth on Cry toxin-incorporated diet in one half of the family should not overlap with QTL for general growth in their siblings reared on a diet lacking Cry toxins. Therefore, larvae from each family were exposed to diets containing either a diagnostic dose of Cry1Ab or Cry1A.105 + Cry2Ab2 expressing corn leaf tissue and leaf tissue from the non-expressing corn near isolines, as described in Dively et al. (2016) and Taylor et al. (2021) (**Figure 1, Figure S1, Table S1, Table S2)**.

**Figure 1.**
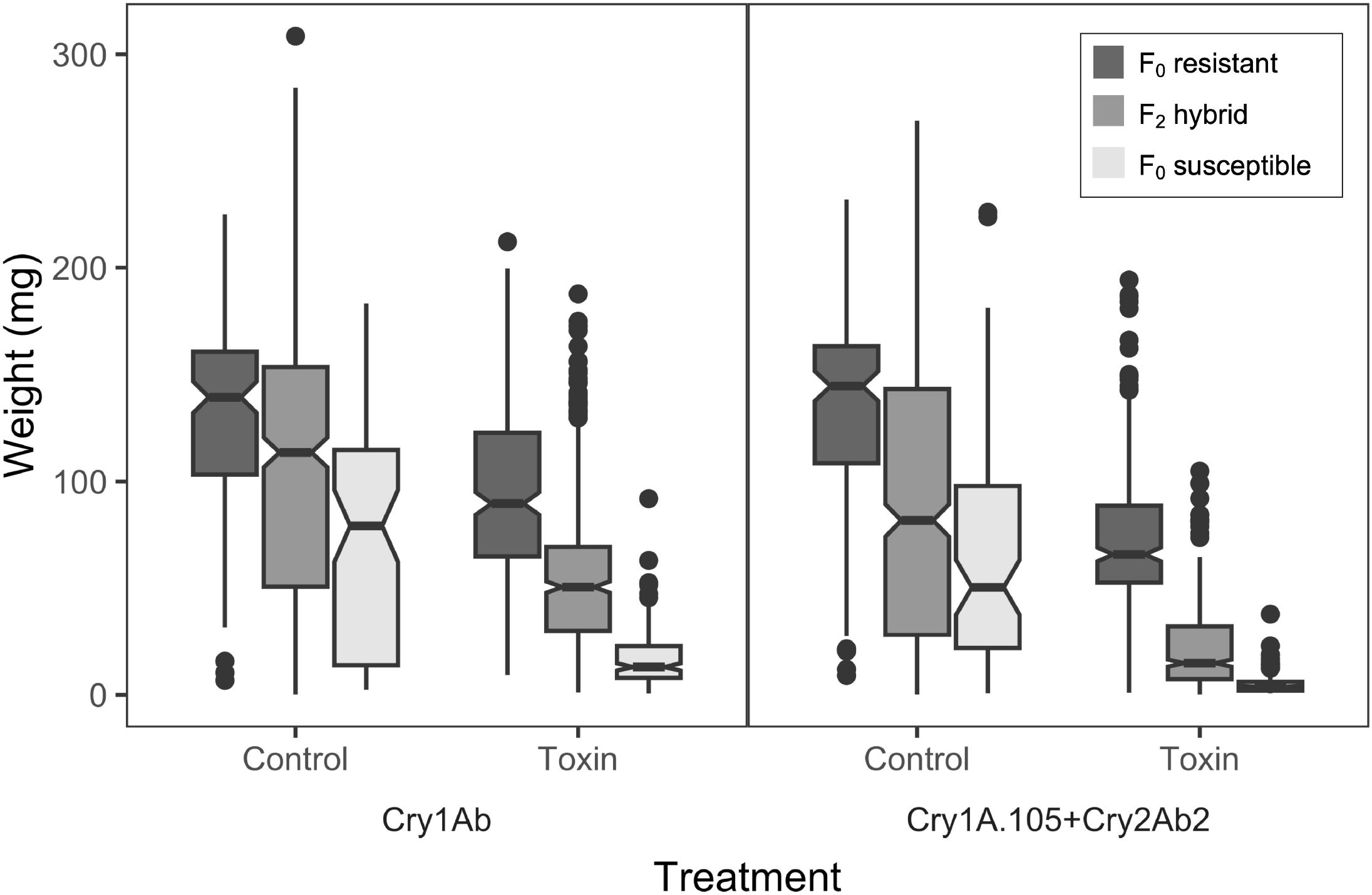
Distribution of weights for *H. zea* individuals after seven days in a laboratory leaf tissue incorporation assay for Cry toxin and control treatments.

The larval population from which resistant cross-founding grandparents were drawn grew significantly larger than laboratory-reared, susceptible larvae on both Cry1Ab and Cry1A.105 + Cry2Ab2 expressing corn leaf tissue incorporated diets **(Figure 1, Table S1)**. Yet even in the absence of Cry toxins, the susceptible population grew significantly less than did the resistant population, suggesting that growth on toxin and general growth were correlated in the grandparental populations. This was likely a result of long-term reproductive isolation between these populations resulting in substantial phenotypic and genomic divergence between them.

Growth phenotypes in the F2 offspring suggested that resistance to all the Cry toxins we tested, as well as growth on the control treatments were quantitative genetic traits (**SI Results**). Based on our experimental design, we predicted that recombination in the F1 generation should reduce correlations between growth on control and toxin-incorporated diet within individual F2 offspring, resulting in limited overlap between general growth and Cry resistance QTLs discovered in our assays.

### Marker generation and variant calling

Grandparents, parents, and progeny from 10 replicated intercross families were sequenced on an Illumina NovaSeq 6000. Whole genome sequencing (WGS) of the 20 cross parents resulted in a total of 4,716,009 high quality genome wide single nucleotide polymorphisms (SNPs) following read filter-trimming and alignment to the *H. zea* genome (v. 1.0, PRJNA767434; Benowitz et al., 2022). Genome wide average sequencing coverage of 18.6⨉ (*st. dev.* = 2.4) ensured accuracy of the genotyping calls. Divergence between the field collected resistant and laboratory susceptible founding populations was high genome wide (**Figures S2 & 3, SI Results**), as would be expected due to the long-term sexual isolation between them.

Double digest restriction site associated DNA (ddRAD) sequencing of F2 offspring resulted in 78,580 - 79,408 high quality SNP markers for genotype-phenotype association in each treatment. Final SNP marker numbers likely varied because of true differences in SNP presence, different loci passing quality control filters, and genetic variation among individuals at restriction cut sites used for marker development (Davey et al., 2013). The average sequencing coverage per ddRAD locus across all F2s was 52,225⨉ (*st. dev. =* 45,011), and the average coverage per locus per individual was 63⨉. SNPs were further filtered to include only those variants where the allele origin population (field resistant vs. lab susceptible) could be reliably predicted to determine their directional effect on growth. This produced the smaller filtered sets of 6,717 - 6,749 genome wide SNPs for each treatment that were used for visualization of effect size and direction.

### Polygenic Cry resistance architecture

To link ddRAD marker genotypes with Cry resistance phenotypes, we used gemma (Zhou et al., 2013). This software estimates both the number of loci underlying complex traits (including those of small effect size), as well as the impact of each large effect locus on a trait, while accounting for the relatedness among individuals in a test population. A significant proportion of phenotypic variation (PVE) in the F2 offspring could be explained by the full set of ddRAD markers. Mean PVE estimates were 69.7% [95% CI = 45.4 - 93.7] for the Cry1Ab treatment and 71.1% [95% CI = 43.6 - 96.5] for the Cry1A.105 + Cry2Ab2 toxin blend, and lower for the control growth treatments 44.7% [95% CI = 20.3 - 71.9] and 51.2% [95% CI = 27.1 - 74.1] (**Figure S4**).

The numbers of variants predicted to have large effect sizes on growth were similar across treatments, with mean estimates ranging from 11.3 - 34.1, further supporting a polygenic trait architecture. In all cases, there were wide overlapping credible intervals around these means, suggesting that tens of genomic variants of major effect underlie phenotypic differences in all growth-related traits (**Figure S4**). Large effect size variants contributed to 74.0% [95% CI = 36.9 - 98.6] of the Cry1Ab growth-related variance explained by our full marker panel (PGE). For the Cry1A.105 + Cry2Ab2 treatment, 77.8% [95% CI = 46.8 - 98.9] of the growth-related variance could be explained by variants of large effect size. These values dropped to 53.4% [95% CI = 0.5 - 97.3%] and 73.6% [95% CI = 24 - 99.2] for the control treatments (**Figure S4**). Notably, the high PVE and PGE estimates for both Cry treatments suggest that our data captured much of the genetic basis of field evolved resistance in *H. zea*.

### Genomic regions underlying Bt Cry resistance

To identify the chromosomal regions underlying field evolved Cry resistance, we estimated the additive effect (β) of a single resistant cross-founder allele on 7 day weight in mg. Smoothed additive effects of genome-wide markers on growth are shown for Cry1Ab, Cry1A.105 + Cry2Ab2, and control treatments in **Figure 2**. The major effect QTL revealed in Figure 2 were also identified using multiple SNP filtering criteria and in the unsmoothed data set (**Figures S5 & 6)**, indicating that our detection of major effect QTL is robust to differences in bioinformatic and analytical approaches. A resistant parent allele in F2 offspring significantly and strongly increased growth on Cry1Ab in regions of Chromosomes (Chr) 2, 3, 6, 9, and 30 with a significance threshold of Bonferroni corrected p < 0.01 (**Figure 2A**). With the less stringent criteria of Bonferroni corrected p < 0.05 we also detected an association of parts of Chr 11 and 21 (**Figure 2A**). For Cry1A.105 + Cry2Ab2, we only detected a significant effect of Chr 9, though non-significant peaks suggest possible shared QTL with the Cry1Ab treatment on Chr 2 and 30 (**Figure 2C**). We detected genomic regions associated with general growth in the laboratory assays on Chr 10, 14 and 27 (**Figure 2B**). As predicted for F2 offspring, we observed little overlap between QTL underlying growth on toxin-containing and control treatments. Chr 10 was the only Chr with a major effect on growth on both control and toxin-containing (Cry1Ab) treatments **(Figures 2A & B)**. Therefore, we did not consider this region to be Cry resistance related. The negative direction of this effect also suggested that the growth associated allele on Chr 10 is more common in the susceptible parent population. Notably, we did not detect any fitness cost of Cry resistance-associated genomic regions, which would have appeared as negative values for these variants in the subsets of F2s on non-Cry expressing treatments (**Figures 2B & D**).

**Figure 2.**
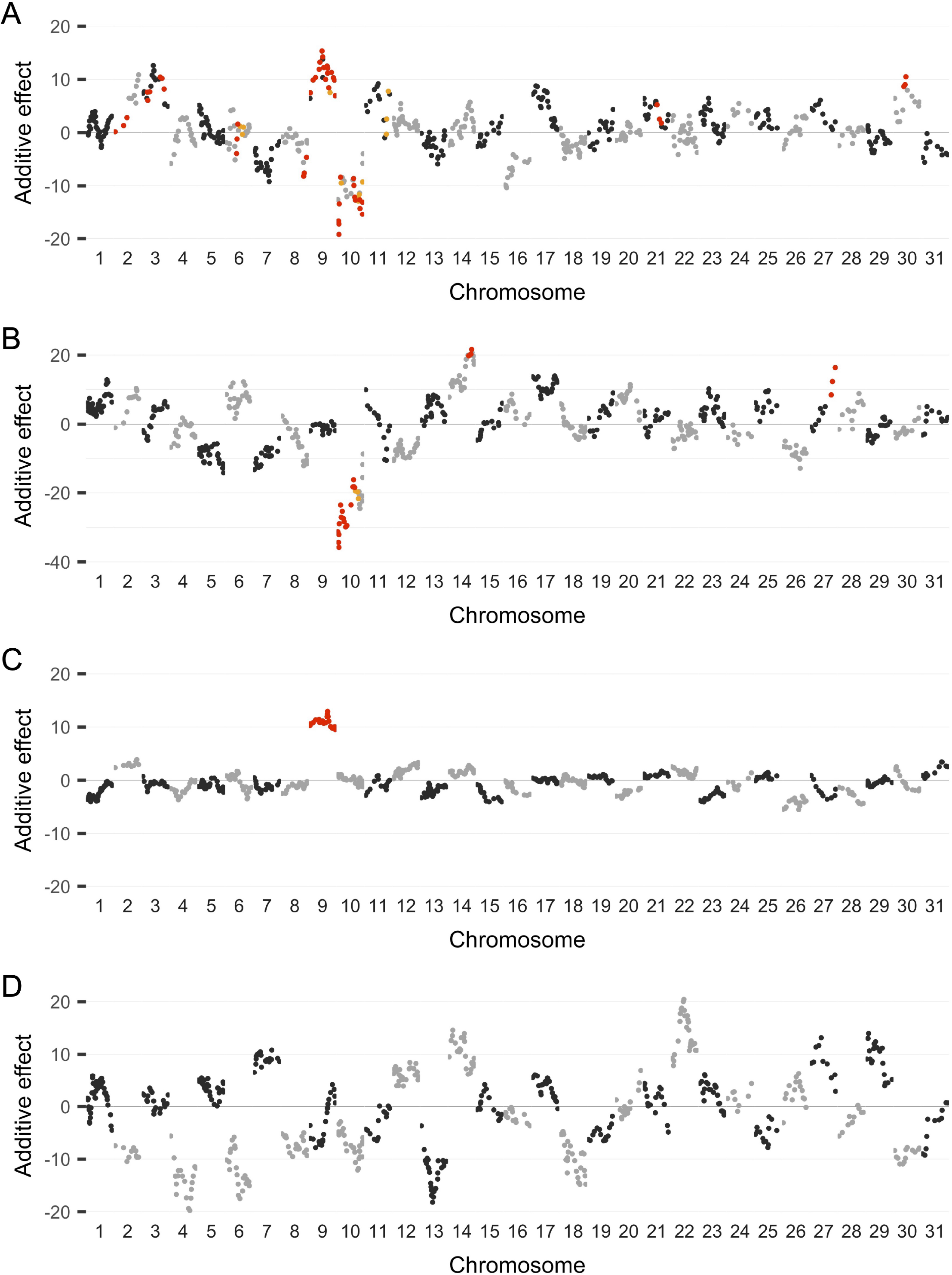
Association of genome wide markers and weight after seven days of feeding on Cry toxin containing and control treatments. The additive effect (beta from LMM) is the weight in mg associated with the presence of a single resistant population allele smoothed for visualization by averaging over 21 SNP windows with a 5 SNP step for markers on 31 *H. zea* chromosomes. The genotype-phenotype association is shown for **A.** weight after seven days of exposure to Cry1Ab, **B.** weight on the non-toxin near isoline control diet for Cry1Ab, **C.** weight after seven days of exposure to Cry1A.105 + Cry2Ab2, **D.** weight on the non-toxin near isoline control diet for Cry1A.105 + Cry2Ab2. Chromosomes are plotted in alternating dark and light gray with each point representing a 21 SNP window. Windows with at least one significantly associated SNP are highlighted in color, with orange indicating an adjusted p value < 0.05 and red indicating an adjusted p value < 0.01.

The region of the genome with the largest additive effect on resistance was on Chr 9 for both Cry-containing treatments **(Figures 2A & C)**. There was no signal of an effect of Chr 9 on general growth, however **(Figures 2B & D)**. For individuals grown on the Cry1Ab treatment, we identified a clear peak between 5 and 6 Mb on Chr 9. This aligned well with previously identified signals of genomic divergence over time at 5.75 Mb on Chr 9 in wild *H. zea* (Taylor et al. 2021). We also identified a directly overlapping divergence peak at 5.75 Mb between resistant and susceptible cross founders (SI Results). Between 5 and 6 Mb on Chr 9, there were 45 genome annotations (**Table S3**), several of which we examined further (see below). The QTL on Chr 30 had clear peaks at ∼2.5 and ∼3.3 Mb, making it possible to describe nearby gene candidates for the Cry resistance observed in our study. Within 100 Kb of those Chr 30 association peaks (2.4 - 3.4 Mb) were a MAP kinase-activated protein kinase 2-like, HzeaTryp129, and HzeaABCC11, toll-like receptors, a cluster of carboxylesterases, and carboxylase-like genes (**Table S4**), suggesting that several known Bt candidate gene families and insecticide resistance related genes could underlie this QTL. The other identified QTL on Chr 2, 3 and 6 had more diffuse genomic signals with less well-defined peaks, making it difficult to identify narrow regions most linked to resistance. In the SI Results, we further describe the limited evidence for any role of other major candidate genes.

### Resistance-associated differential gene expression

We paired our QTL study with differential gene expression analyses to narrow in on potential targets of selection. The mechanism of Cry toxin action takes place in the larval midgut, and changes in gene expression in this tissue have strong potential to impact resistance. Therefore, we initially focused on differential midgut gene expression between wild, resistant *H. zea* larvae collected directly from Cry1A.105+Cry2Ab2-expressing corn and laboratory-reared, susceptible larvae. Of the 14,600 annotated genes, 527 genes were significantly upregulated and 352 were significantly downregulated in the population collected from Cry expressing corn relative to the susceptible population (p-adjusted < 0.01; **Table S5**). Of those significantly differentially expressed genes, 33 were within 100 kb of at least one SNP significantly associated with resistance to Cry1Ab, Cry1A.105 + Cry2Ab2, or both treatments (**Table S6**). This analysis identified several significantly upregulated genes in resistant larvae that were found near QTL, signaling their potential involvement in resistance. One was an aminopeptidase gene found on Chr 9 (*apn1*) (**Table S6**). *apn1* is a known Cry resistance candidate gene, for which reduction of expression increases Cry resistance (Herrero et al., 2005; X. Ma et al., 2022; Sun et al., 2022). In our work, however, *apn1* expression increased in our resistant population, which was inconsistent with its previously described role in resistance. The most striking gene expression differences were found among seven significantly upregulated trypsin genes arranged in tandem between 5 and 6 Mb in the Chr 9 QTL. Six of these trypsins in the tandem array on Chr 9 were among the top 50 most differentially expressed genes genome wide **(Figure 3B, Table S5)**.

**Figure 3.**
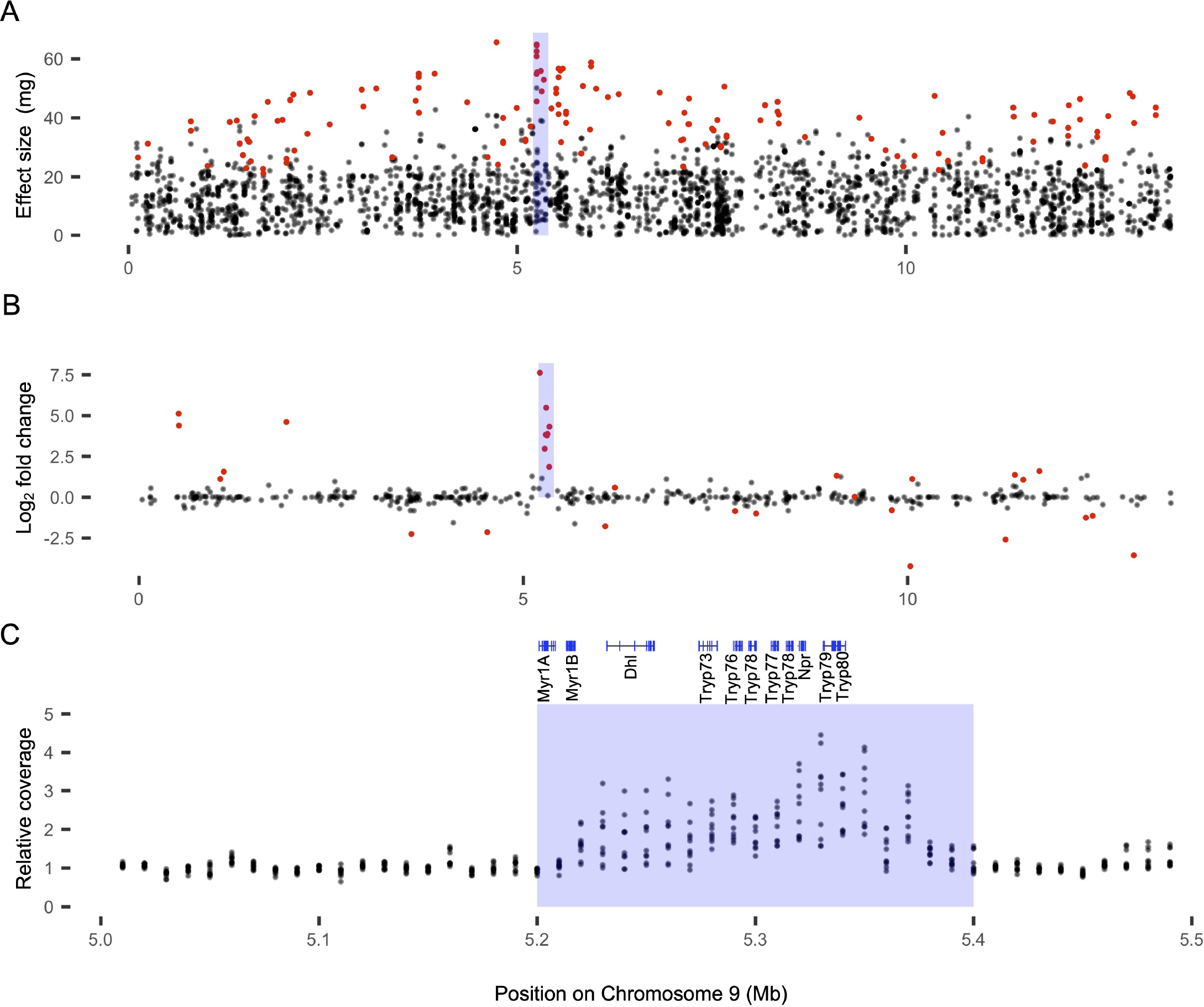
Genomic signals indicate that a region between 5.2 and 5.4 Mb on *H. zea* Chr 9 is duplicated in field resistant individuals and associated with resistance phenotype. The absolute value of the effect size (beta from the LMM) for ddRAD SNPs is plotted in (**A**). A differential expression plot for all genes on Chr 9 between resistant and susceptible populations is shown in (**B**). In (**C**) the gene annotations between 5.2 and 5.4 Mb on Chr 9 are shown above a plot of relative depth of sequencing coverage for field resistant *H. zea*. Gene abbreviations are as follows: Myr1 = myrosinase 1-like, Dlh = disks large homolog 4-like, Tryp = Trypsin, Npr = neuropeptide receptor A35. In all panels the region between 5.2 and 5.4 Mb on Chr 9 is highlighted in blue.

Trans-acting factors also impact regulation of gene expression, which would result in separation of differentially expressed genes from QTL. When we considered only our differential gene expression analysis, six other trypsins spread across multiple chromosomes were also among the top 50 most differentially expressed genes (**Table S5**). Several other insecticide resistance candidate gene families also had members among the top 50 most differentially expressed genes, including, a cadherin-87A-like gene and a cytochrome P450 (CYP301B1), both of which were downregulated in resistant individuals (**Table S5**).

We also tested for Cry inducible gene expression changes between wild *H. zea* larvae collected from Cry expressing and non-expressing corn. A comparison of midgut gene expression for these groups revealed that expression of a small number of genes was modulated by Cry toxin exposure. Thirteen genes were significantly upregulated and 20 genes significantly downregulated in larvae collected from expressing corn (**Table S7**). Four trypsins and one chymotrypsin were among the most upregulated genes in Cry exposed resistant individuals. Only one of the trypsins found between 5 and 6 Mb on Chr 9, *tryp80*, was in this group showing Cry inducible expression changes. Downregulated genes in Cry exposed individuals included a cadherin-87A-like gene and the immune-related gene, Lysozyme1.

### A potential genetic mechanism of large effect on Bt Cry resistance

Taken together, evidence from the QTL and differential expression analysis suggested that a region of Chr 9 containing a cluster of differentially expressed genes, including 7 trypsins, was strongly associated with Cry resistance **(Figures 3B & C)**. Closer investigation of this region revealed a structural variant from 5.22 - 5.37 Mb on Chr 9 **(Figure 4A)**. In this region encompassing the full cluster of differentially expressed genes, resistant cross founders had increased whole genome sequencing depth of coverage relative to their genome wide average **(Figure 4A)**. Elevated coverage depth indicated this region of the genome was at least duplicated in resistant cross founders. The observed upregulation of most genes in this gene cluster in resistant individuals was consistent with a mutational event that duplicated the entire gene cluster **(Figure 3B)**.

**Figure 4.**
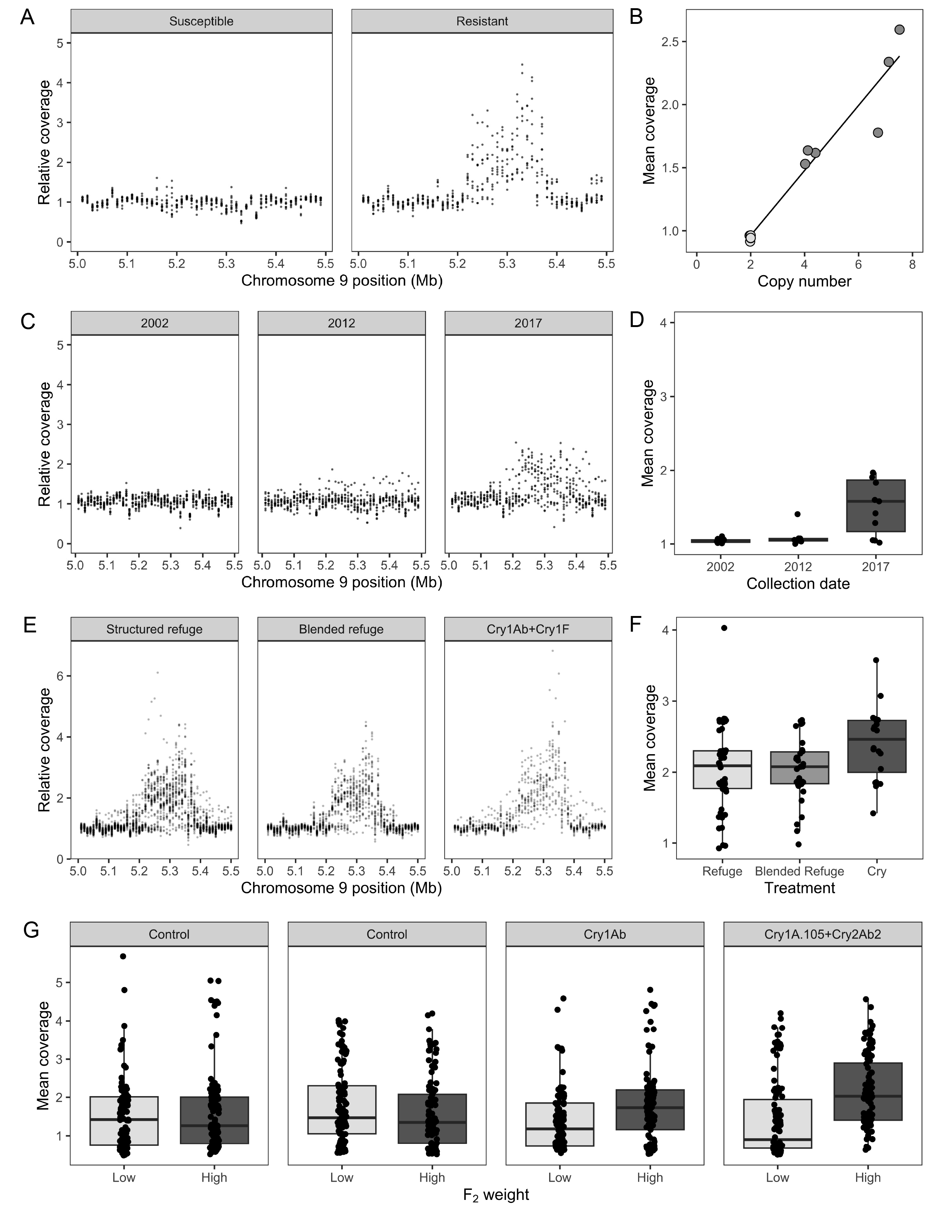
Resistance evolution and the gene cluster amplification on *H. zea* Chr 9. **(A)** Relative sequencing depth is plotted for susceptible and resistant cross founders. In A, C, and E each point represents the average sequencing depth in that 10 kb window for a single individual relative to that individual’s genome wide average sequencing coverage. **(B)** Coverage is associated with ddPCR copy number variant analysis for *trypsin 77* (r^2^ = 0.95) for the resistant (dark gray) and susceptible (light gray) founders of mapping families. **(C)** Relative sequencing depth is plotted for individuals collected in LA in 2002 before Cry resistance evolution, 2012 as resistance was evolving, and 2017 after practical resistance was first detected. **(D)** The mean coverage depth of the amplified region from 5.22 - 5.37 Mb is plotted for the same individuals shown in C. **(E)** Relative sequencing depth is plotted for individuals collected in 2019 in NC from structured refuge, blended refuge and Cry1Ab+Cry1F expressing corn. **(F)** The mean coverage depth of the amplified region is plotted for the same individuals shown in E. Coverage was significantly higher for the individuals taken from Cry expressing plants compared to both refuge types (t = 2.61, p = 0.039; t = 2.67, p = 0.033), suggesting selection for higher copy numbers. **(G)** Mean ddRAD sequencing coverage for F2 offspring that were the top and bottom for 7 day weight in each treatment. Relative coverage was significantly higher for the F2 offspring that grew largest on both toxins (t = 3.46, p < 0.001; t = 5.21, p < 0.001). On both control treatments there was no relationship between coverage and growth (t = 0.46, p > 0.05; t = -1.51, p > 0.05).

To confirm that the coverage and differential expression signals we detected were truly related to gene cluster duplication, we quantified the copy number variation for one representative gene in the cluster (*tryp77*) using droplet digital polymerase chain reaction (ddPCR) **(Figure 4B)**. If *tryp77* existed as a single copy gene, our ddPCR results should reveal a copy number equal to two in this diploid species. Instead, our analysis showed that all resistant cross founders from our study had more than two copies of *tryp77*, while all susceptible founders had the expected two copies. Variation in copy number existed among the field-collected resistant individuals used in our analysis, indicating there is not a single allele, but multiple alleles linked to resistance at this Chr 9 locus. There was a strong relationship between copy number detected by ddPCR and relative coverage depth of the gene cluster (r^2^ = 0.95), suggesting that WGS coverage depth is a reasonable proxy measure for copy number.

We used publicly available WGS data from Taylor et al. (2021) and Pezzini et al. (2024) to determine whether variation in copy number at this Chr 9 locus could also be found in other wild North American *H. zea* populations. Increased depth of coverage was not found in WGS data from the susceptible lab population or in field samples collected in 2002, decades before Cry resistance was widespread **(Figures 4A-D)**. Absence of this variant in samples collected in 2002 suggested that: 1) it may have arisen after 2002, 2) it existed as a rare allele at our study site prior to 2002, or 3) it arose in or before 2002, but at a location not sampled for our work.

Analysis of WGS data from these *H. zea* samples collected from Louisiana in 2017 **(Figures 4C & D)**, and North Carolina in 2019 **(Figures 4E & F)** revealed signals of increased coverage depth in this Chr 9 region, similar to what we observed in the resistant cross founding individuals collected from Maryland in 2019 and 2020 **(Figure 4A)**. The presence of this sequence duplication in samples collected across 3 separate regions of North America over multiple years, suggests that this genetic variant is widespread. Notably, depth of coverage at this locus increased over time, with increased coverage depth first appearing in 2012 and the strength of that signal increasing in 2017 **(Figures 4C & D)**.

In 109 samples collected from experimental plots in North Carolina in 2019, we detected increased coverage depth in this Cry resistance-associated Chr 9 region in individuals collected from Cry expressing plants compared to individuals collected from non-Bt plants in structured and blended refuge plots (Structured refuge: t = 2.61, adjusted p = 0.039, Blended refuge: t = 2.67, adjusted p = 0.033) **(Figure 4E & F)**. The consistent difference in coverage depth between the population collected on Cry expressing plants and both refuge conditions and suggests that selection by the Cry expressing plants limited the growth and survival of individuals with lower sequence duplication in this region, providing another line of evidence for the role of this genetic variant in Cry resistance evolution. In these samples, we also observed a continued increase in sequencing depth over time, with samples from 2019 and 2020 (**Figure 4A & E**) showing stronger signals of this amplification than those from 2012 and 2017 (**Figure 4C**). This could reflect increased numbers of individuals homozygous for a duplication or additional duplication events. Though none of the resistant samples collected in MD and NC in 2019 and 2020 appear to be single copy in this region (**Figure 4A & E & F**), some individuals collected from non-expressing plants are (**Figure 4E & F**), indicating that this variant is likely at high frequency but not fixed in modern populations. Samples collected in 2019 and 2020 also have markedly higher coverage between 5.32 and 5.35 Mb, region contains *tryp79* and *tryp80,* relative to other parts of the duplicated cluster and historical samples. We speculate that this particular region may have undergone multiple duplication events.

The strong association of *tryp77* copy number and depth of WGS coverage motivated us to analyze the correlation between ddRAD sequencing coverage depth and weight for the F2 offspring from our experimental crosses **(Figure 4G)**. We reasoned that F2 individuals with higher coverage at this locus should, on average, have higher weights than those with low coverage, if copy number variation was involved in Cry resistance. Likewise, if increasing copy number is only related to Cry resistance and not generally associated with weight gain, we reasoned that any positive correlation between weight and coverage depth should be observed for individuals fed on Cry-treated diet, but not for their siblings on untreated diet. Indeed, ddRAD- seq coverage depth of the Chr 9 gene cluster was significantly higher for the F2 offspring that grew most on both toxin containing diets compared to the offspring which grew the least (Cry1A.105 + Cry2Ab2: t = 5.2054, p adjusted < 0.001, Cry1Ab: t = 3.4574, p adjusted <0.001) **(Figure 4G)**. On both control diets, there was no relationship between relative coverage and growth (Control 1: t = -1.5115, p = 0.1323, Control 2: t = 0.46397, p = 0.6432) **(Figure 4G)**. As this is a polygenic trait and the Chr 9 locus was not the only one to confer Cry resistance **(Figure 2A)**, we expected that F2 genotype at this locus would not perfectly predict phenotypes.

However, the strong association of ddRAD-seq coverage with resistance phenotype indicates that the region containing this amplified gene cluster likely explains most of the association between Chr 9 and Cry resistance.

### Synergistic effect of trypsin inhibition and a Cry toxin

Ten genes were found in the region of copy number variation on Chr 9, and seven of these genes encoded trypsins. To assess whether trypsins, including those on Chr 9, were involved in Cry resistance, we inhibited their activity with N-α-tosyl-ւ-lysine chloromethyl ketone (TLCK) in resistant and susceptible populations of *H. zea*. If trypsin activity, including the activity of those on Chr 9, were involved in Cry resistance, we reasoned that their inhibition should interfere with growth on Cry treated diet. Trypsins are generally involved in lepidopteran metabolism (Muhlia-Almazán et al., 2008), and their inhibition could also lead to stunted growth on untreated diet. Yet if the midgut expressed trypsins on Chr 9 had activity unique to Cry resistance, we reasoned that increasing dosages of a trypsin inhibitor should more strongly impact the weights of larvae feeding on Cry-treated diet, compared to those fed trypsin inhibitor on an untreated control diet. This trend should be true for both susceptible individuals bearing single copy genes in the cluster, as well as resistant individuals with copy number variation. We solubilized TLCK in a phosphate saline buffer (PBS) to make different concentrations of trypsin inhibitor, and each concentration was mixed with our leaf tissue-incorporated diets. Early second instar larvae were grown on either a diagnostic diet of Cry1Ab expressing corn leaf tissue or leaf tissue from the non-expressing near isoline, each with increasing dosages of TLCK. Larval weight was examined after a 7 day diet exposure. We confirmed that incorporation of PBS did not impact larval growth relative to our standard non-expressing leaf tissue diet (p > 0.05, data not shown), and therefore larval growth on buffer alone was used as a control for all further analyses and visualization.

To understand the extent to which trypsin inhibition influenced Cry resistance, we calculated a growth ratio, which compared growth on the Cry-treated and untreated diets at each TLCK dose. If trypsins, including those in our Chr 9 cluster, showed no special activity related to Cry resistance, we reasoned that the ratio of larval weights on these diet treatments should remain constant with increasing doses of a trypsin inhibitor. Instead the growth ratio was negatively correlated with TLCK dose, suggesting that increased suppression of trypsin activity had a proportionally greater impact on the weights of larvae grown on Cry-treated diets, than those on untreated diet **(Figure 5)**. The interaction between TLCK dose and Cry1Ab exposure on larval weight was statistically significant for both resistant and susceptible *H. zea* populations (p < 0.001) (**Figure 5, Table S8**), and post hoc contrasts of the estimated marginal means confirmed that a trypsin inhibitor paired with Cry1Ab toxin significantly decreased larval growth (**Table S9**). Notably, the mean weight of larvae from the resistant population fed on the Cry1Ab and 10X TLCK treatment was 17.4 mg, suggesting that strongly inhibiting trypsins in resistant individuals could reduce their Cry tolerance to levels observed in the susceptible population (mean = 20.7 mg; **Table S8)**. These results failed to falsify a role for trypsins in Cry resistance, including those from our Chr 9 copy number variant (CNV). Overall, our findings suggest that trypsin activity serves as one protective mechanism against Cry1Ab toxin expressed in corn.

**Figure 5.**
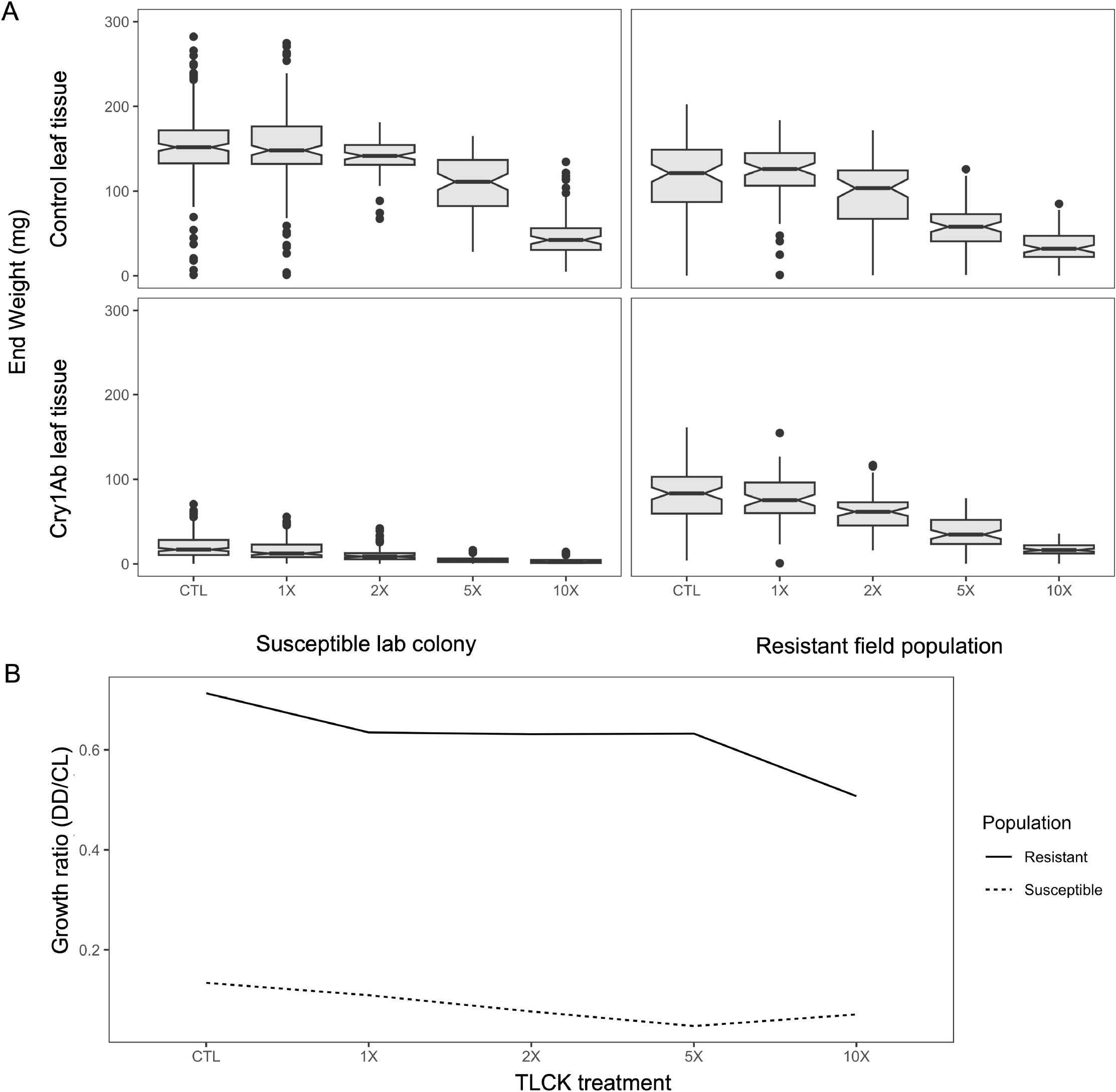
Effect of trypsin inhibition. **(A)** Distribution of weights for laboratory susceptible and field resistant *H. zea* individuals after seven days in a trypsin inhibition assay with Cry1Ab containing corn leaf tissue and leaf tissue from the non-expressing near isoline. **(B)** The growth ratio for each trypsin inhibitor dose, a measure of how much weight is suppressed by Cry toxins and TLCK combined compared to the effects TLCK alone.

## Discussion

Here, we confirm the polygenic nature of *H. zea*’s adaptation to transgenic crops in agroecosystems, and we begin to shed light on the mechanisms underlying their rapid evolutionary response. Using replicated, split-family QTL analyses, we identified multiple genomic regions associated with Bt resistance, most of which are not shared by siblings that grew well on untreated diets (**Figure 2**). This lack of overlap demonstrates that the multiple genomic regions associated with growth on Cry-containing diets are Cry resistance loci.

Polygenic adaptation to Cry toxins was predicted for *H. zea* because selection by Bt crops was expected to act within its broad phenotypic range of Cry tolerance values. Early studies of field-collected *H. zea* described this range and documented that laboratory selection could rapidly increase population-level Cry tolerance (Luttrell et al., 1999). Our data provide empirical support for the prediction that polygenic adaptation is favored when selection acts on phenotypes within the natural range of variation of a population (McKenzie and Batterham 1994).

Tens of loci contribute to the polygenic basis of Cry resistance in *H. zea*, but the genomic region of largest effect on both Cry1Ab and Cry1A.105 + Cry2Ab2 resistance was found on Chr 9. All resistance associated genomic regions are likely to be important to the Cry tolerance observed in wild *H. zea*, and other loci should be the focus of future investigations. It is likely that genetic interactions among the Cry resistance loci revealed in our work are critical for production of strong Cry resistance phenotypes, as is observed in other species (Ma et al. 2022,

Sun et al. 2022). However, much of our analyses focused on Chr 9, providing us with insight into the nature of the genetic mutations that facilitated rapid evolution in this system. We used multiple lines of evidence to document the nature of the Chr 9 locus and demonstrate its importance to Cry1Ab resistance both in our lab-based QTL experiments, as well as in field-collected populations.

For our first line of evidence, we showed that field-collected Cry resistant *H. zea* overexpressed multiple genes in a 10 gene cluster situated within the Chr 9 genotype-phenotype association peak (**Figure 3**). Changes in gene expression are well known to underlie adaptive phenotypes, including for cases of rapid insecticide resistance evolution (Amezian et al., 2021; Guo et al., 2021; Nauen et al., 2022; Wilding, 2018). An alternative explanation for increased expression of an entire gene cluster is sequence duplication, however (Heckel, 2022; Kondrashov, 2012). We used ddPCR of our cross-founding parents to demonstrate that field-evolved Cry resistant *H. zea* currently carry multiple copies of at least one representative gene (*tryp77*) in the cluster. From this, we concluded that a CNV found in the 5-6 Mb region of Chr 9 was strongly (but not exclusively) associated with field-evolved Cry resistance in *H. zea*. It was not possible to reconstruct the full sequence of this large CNV with the short read sequencing approaches used in this study, and future studies should employ long read sequencing to describe haplotypic variation in this genomic region.

Our second line of evidence for the role of this region in Cry resistance was the change in copy number variation on Chr 9 over time in wild *H. zea*. Changes in Cry1Ab and Cry1A.105+Cry2Ab2 tolerance in wild *H. zea* slowly increased following commercialization of Bt crops in 1996 (Dively et al. 2016). Early warning signs of resistance emerged in 2008 (Brévault et al., 2013) and widespread practical resistance to both Cry1A and Cry2A toxins was documented by 2016 (Dively et al., 2016; Reisig et al., 2018; US-EPA, 2018; Yang et al., 2019). Using WGS read coverage depth as a proxy for copy number, we documented an increase in copy number at this locus in wild *H. zea* over time. Consistent with previous resistance observations, *H. zea* collected from LA in 2002 showed no evidence of a CNV, those from 2012 showed evidence of a rare CNV, and most 2017 individuals carried the variant. This indicated that the Chr 9 CNV was widespread by the time *H. zea* could readily feed on Cry-expressing corn ears in the field.

Our recent collections of the cross-founding parents from MD in 2019 and 2020 showed that the Chr 9 CNV was not fixed for a specific copy number. While all resistant cross founders carried a Chr 9 CNV, some carried different numbers of copies **(Figure 4B)**, which likely confer different levels of fitness under selection pressure by Cry toxins. This idea motivated the analysis which provides a third line of evidence for the involvement of the Chr 9 CNV in *H. zea*’s rapid adaptation to Bt crops. A re-analysis of WGS data from wild *H. zea* collected as part of a field experiment in NC in 2019-2020 (Pezzini et al. 2024) documented that the Chr 9 locus continues to be under strong selection by Cry toxins. Larvae collected directly from Cry1Ab+Cry1F expressing field corn showed significantly greater WGS coverage depth of this loci, our proxy for copy number variation, than did individuals collected from structured or blended refuge ears **(Figures 4E & F)**. We previously showed that SNPs in this CNV-containing region were under strong Cry selection (Pezzini et al. 2024), but here we demonstrated that a generation of selection by Cry1Ab+Cry1F increased the copy number in wild *H. zea*. Moreover, coverage signals suggested that no individuals collected from Cry1Ab+Cry1F expressing corn had single copy genotypes, while their conspecifics collected from non-expressing corn in the same small experimental plots did. These data suggest that the CNV on Chr 9 continues to be under selection for increased copy number, even though copy number variation is widespread in the North American landscape.

Interestingly, seven of the ten genes within the CNV were trypsins, all of which were differentially expressed **(Figure 3B)**. The highest WGS coverage depth in wild *H. zea* collected on Cry1Ab+Cry1F expressing corn in NC also spanned two of those trypsins. Moreover, expression of one of the trypsin genes within this cluster, *tryp80*, appeared to be inducible upon exposure to Cry-expressing corn **(Table S7)**. Trypsins are a gene family previously thought to be involved in lepidopteran Cry resistance via toxin activation or degradation (Gong et al., 2020; González-Cabrera et al., 2013; C. Liu et al., 2014; Y. Ma et al., 2013). We demonstrated that trypsin inhibition synergistically reduced growth in Cry containing treatments for both resistant and susceptible populations. These findings demonstrated a significant protective effect of trypsin activity in Cry exposed *H. zea*. When exposed to the highest TLCK dose and Cry toxin together, growth in the resistant population was suppressed to the level of the susceptible population on Cry toxin alone. The shared response of susceptible and resistant populations to TLCK in this assay suggests that the increased expression and duplication of trypsins might allow the resistant population to better utilize a mechanism already present in the susceptible population.

Previous work has also linked trypsin activity to Cry resistance in *H. zea*, for example, Lawrie et al. (2020) found strong signals of trypsin upregulation in resistant *H. zea* while Zhang et al. (2019) found that trypsin down regulation was related to *H. zea* Cry resistance in a laboratory selected line. The upregulation of trypsins in resistant individuals observed here suggests degradation rather than activation as the potential mechanism for these genes’ involvement in resistance. This would be expected in the case of resistance to crops that express activated toxin. Phenotypic resistance due to Cry toxin degradation by trypsins has been reported in field populations of *Spodoptera exigua* (Y. Ma et al., 2013). Our findings along with results from other Lepidoptera suggest a potential resistance mechanism: gene amplification causes increased expression of these trypsins which, in turn, may degrade activated Cry toxins.

The CNV on Chr 9, likely plays an important role in resistance, but we emphasize that it is not the only region involved in *H. zea*’s rapid adaptation to Bt crops. We also identified resistance associated genomic regions on Chr 2, 3, 6, and 30. Multiple promising candidate genes were found in the QTL peak on Chr 30 and near other QTL. There were also multiple candidate Bt resistance genes among the top differentially expressed genes between resistant and susceptible *H. zea,* including a cadherin gene. The genomic architecture of Bt resistance likely involves interactions between some of these candidates of large to moderate effect size, as well as undetected small effect size loci to produce the resistance observed in wild *H. zea*. Indeed, our results showed that, on average, large Cry-exposed F2s had significantly higher depth of sequencing coverage, and therefore copy number in our Chr 9 CNV, than did smaller Cry- exposed F2s (**Figure 4G**). However, some small F2s contained the Chr 9 CNV, while some large F2s did not. As expected based on the multiple QTL regions uncovered in our analysis, this suggests one or more additional segregating resistance alleles are additively or synergistically interacting with the Chr 9 CNV to produce resistance phenotypes. Consistent with our findings, interactions between mutations in multiple genes have been described in laboratory studies of other lepidopteran species (Ma et al. 2022, Sun et al. 2022). Further experimental work targeting the Cry resistance associated regions on Chr 2, 3, 6 and 30 will be necessary to identify the additional mutations underlying field-evolved Cry resistance.

In our non-expressing control growth treatments, we did not detect any fitness costs of this Cry resistance associated gene amplification, or any of the other detected major effect loci (**Figure 2**). The lack of fitness cost for resistance, the low dose of the toxins, incomplete refuge implementation, and cross pollination, likely contributed together to the failure of Cry resistance management strategies for *H. zea*. A degradation mechanism for toxin resistance could explain the rapid evolution of resistance to multiple toxins that do not share binding sites. Our findings link the same gene amplification to resistance to both Cry1Ab and blends of Cry1A.105 + Cry2Ab2 and Cry1Ab+Cry1F. It may also be associated with resistance to other Cry toxins not tested here. If, as our results suggest, trypsin activity underlies Bt resistance phenotypes in field *H. zea* populations, then targeted synergist trypsin inhibitors could potentially be developed to restore efficacy of Bt crops in controlling *H. zea* (Correy et al., 2019).

Gene duplications, an important source of genetic variation for phenotypic evolution, have been linked to many cases of insecticide and herbicide resistance evolution (Bass & Field, 2011; Heckel, 2022). Duplications of detoxification related genes can immediately confer resistance by increasing the quantity of already functional detoxification enzymes or can additionally lead to new mechanisms of resistance by neofunctionalization. Multiple cases of insecticide resistance have been functionally linked to new duplications of cytochrome P450s and carboxylesterases, two detoxification related gene families (Bass et al., 2013; Cattel et al., 2021; Mouches et al., 1990; Puinean et al., 2010; Schmidt et al., 2010). Our findings here suggest a new example of the duplication of a detoxification related gene leading to insecticide resistance, and the first case of this mechanism linked to Bt resistance. It is possible that gene duplication may be more likely to underlie rapid phenotypic evolution than other variant types, but the prevalence of structural variants is not well described as detection is difficult with current technologies (Ho et al., 2020; Mahmoud et al., 2019). As gene amplifications have frequently been linked to resistance evolution, genomic monitoring for resistance should be extended to consider a range of genomic variant types beyond SNPs (Fritz 2022).

## Methods

### Insect samples and phenotypes

Field samples of Cry resistant *H. zea*, defined as surviving to late instar on a Bt corn plant expressing Cry1A.105 + Cry2Ab2, were collected in 2019 and 2020 in Prince George’s County (MD) at the University of Maryland CMREC farm in Beltsville. Late instars were collected from Cry1A.105 + Cry2Ab2 expressing sweet corn and reared in the lab on a 16:8 long day light cycle at 25°C and 50% room humidity. Newly emerged adults from field collections were single pair mated to produce a second generation of resistant field derived *H. zea*. Susceptible *H. zea* were acquired from a population that has been maintained in the laboratory without exposure to Cry toxins at Benzon Research (Cumberland County, PA). Cry tolerance in both the resistant field and susceptible lab populations was measured with a diagnostic dose in a corn leaf tissue incorporation assay as described in Taylor et al. (2021) to confirm resistant and susceptible status. Briefly, early second instar caterpillars were fed for seven days on a laboratory diet mixed with powdered lyophilized leaf tissue from Bt expressing corn and their non-Bt expressing near isolines. Resistance was measured by weight gain after seven days on Bt expressing leaf diets.

Weight gain on the non-expressing near isoline control diets served as a measure of non-resistance associated growth.

The field derived progeny that grew well in the Cry expressing resistance assay (not in the bottom quartile for growth) were single pair mated to a susceptible individual from the lab colony. The specific susceptible individuals used in crosses were not exposed to the Cry toxin resistance assay due to the significant growth suppression that would be experienced by this highly susceptible population. Offspring of the single pair matings between the resistant and susceptible populations were sibling mated to produce an F2 generation. Cry toxin resistance phenotypes were assessed in the F2 progeny also using the leaf tissue incorporation assay, as described in Taylor et al. (2021). Five F2 intercross families were assessed for resistance to Cry1Ab by measuring growth on diet with incorporated powdered Cry1Ab expressing leaf tissue. The other half of the offspring were assayed for growth related advantages on the diets with powdered leaf tissue from the non-expressing near isoline of Cry1Ab expressing corn. Five different intercross families were assessed for resistance to Cry1A.105 + Cry2Ab2 by measuring growth on diet with leaf tissue expressing both toxins for half of the offspring. The other half of the offspring were used to assess growth related advantages on diets containing leaf tissue from the non-expressing near isoline. Between 146 and 282 F2 offspring were split across treatments and phenotyped from each family, resulting in 73 to 142 F2 offspring phenotyped per treatment from each family. F2 numbers varied due to differences in larval availability for assays. After phenotyping, the larvae were grown on a non-expressing lab diet to increase in size before DNA isolation. All samples were flash frozen and stored at -80℃.

The impact of experimental treatment and population on weight phenotypes after seven days of growth in the laboratory assays was assessed using a model reduction approach between nested general linear models all with a gamma distribution, as is appropriate for the continuous and positive weight phenotypes (Bolker, 2008). The treatments compared were 1) Cry1Ab expressing corn leaf tissue, 2) control tissue from the non-expressing near isoline of Cry1Ab corn, 3) Cry1A.105 + Cry2Ab2 expressing corn leaf tissue, and 4) control tissue from the non-expressing near isoline for Cry1A.105 + Cry2Ab2 corn. The populations compared were 1) the resistant field derived F0, 2) susceptible lab colony F0, and 3) the intercross F2 offspring. The models were compared using a likelihood ratio ꭓ^2^ test with an *a priori* α of 0.01. Post hoc pairwise contrasts were calculated with the R package emmeans (v. 1.8.6; Lenth et al., 2023) with a bonferroni correction to p-values.

### DNA extraction and sequencing

Individuals in the top and bottom 20% of weight phenotypes from each treatment and family were selected for DNA extraction and sequencing. When individuals in the top or bottom 20% did not survive to preservation, they were replaced by individuals in the top or bottom 30%. Genomic DNA was extracted from 833 high and low weight intercross F2 offspring and all F1 and F0 parents with a DNeasy (Qiagen, Hilden, Germany) extraction kit using the modified mouse tail protocol of Fritz et al. (2020). Larval and pupal tissue from the rear ½ - ⅓ of the insect was used for extraction, while for adults, ½ of the thorax was used. DdRAD libraries were prepared for all F2 samples as described in Fritz et al. (2016, 2018). Briefly, samples were digested with the restriction enzymes EcoRI and MSPI (New England Biolabs, Ipswich, MA), and unique 6-mer or 8-mer barcodes were annealed prior to pooling with 38 - 39 other samples. Pools were size-selected for 450-650 bp fragments using a Pippin Prep (Sage Scientific, Beverly, MA), then 12 replicated PCR reactions with 14 cycles each were used to amplify DNA and add a standard Illumina TruSeq index as an identifier to each pool. Pools were sequenced at North Carolina State Genomic Sciences Laboratory on an S4 flow cell of the Illumina NovaSeq 6000, resulting in 173.9 - 97.5 million raw 150 bp paired-end reads per sample pool with an average of 128 million reads per pool. Additionally whole genome sequencing was performed for the 20 cross founding parents on part of an S4 flow cell of an Illumina NovaSeq 6000 at the University of Maryland Baltimore Institute for Genome Science. This sequencing resulted in 30.9 - 48 million raw reads per sample with an average of 39.5 million raw reads.

### Bioinformatic analysis

F2 ddRAD sequencing reads were demultiplexed and quality controlled using the stacks script process_radtags (v.2.61; Rochette et al., 2019). The reads were only retained if they had exact matches to the correct barcode sequence, quality scores above 20 in all 15 bp length sliding windows, and no evidence of adapter contamination allowing up to 2 bp mismatch in adapter sequence. Quality controlled reads were aligned to a *H. zea* chromosome scale assembly (v. 1.0, PRJNA767434; Benowitz et al., 2022) with Bowtie (v.2.2.5; Langmead & Salzberg, 2012) using the “very sensitive” alignment option. Variants were called with bcftools mpileup (v. 1.9; Danecek et al., 2016) and pruned to include biallelic SNPs with a minor allele frequency > 0.05, with a quality score > 50, present in > 50% of samples. The average sequencing coverage per individual at a called ddRAD locus in the final variant set was 63⨉. The total number of ddRAD sequencing markers remaining for genotype phenotype association ranged from 78,580 - 79,408 (Cry1Ab treatment group = 79,261, Cry1Ab control group = 79,182, Cry1A.105 + Cry2Ab2 treatment group = 79,408, Cry1A.105 + Cry2Ab2 control group = 78,580).

Whole genome sequencing reads were quality controlled with Trimmomatic (v. 0.39; Bolgar et al 2014), to remove Illumina adapters, low quality regions at the beginning or end of the read or where the average quality of a four base pair window fell below 30, and any reads shorter than 36 bp. Trimmed reads were aligned and variants were called as described above for the ddRADseq reads. In the filtered WGS data, a total of 4,716,009 high quality SNPs were identified between resistant and susceptible populations, with a genome wide average sequencing coverage depth of 18.6⨉ (*st. dev.* = 2.4). For those analyses of the ddRAD sequencing data that identified the direction of allele effect, SNPs were further filtered to include only those variants where the allele origin (resistant or susceptible population) could be predicted with some certainty. To identify SNPs where the likely origin population of the allele could be predicted, only those SNPs with an allele count difference > 15 (out of a possible 20 alleles in each population) between the resistant and susceptible parent populations were used. The population where the allele was more common was used to identify the allele as resistance or susceptibility associated. After filtering for effect direction, 6,717 - 6,749 SNPs remained (Cry1Ab treatment group = 6,737, Cry1Ab control group = 6,749, Cry1A.105 + Cry2Ab2 treatment group = 6,733, Cry1A.105 + Cry2Ab2 control group = 6,717).

### Resistant and susceptible founding populations genome wide comparison

Genome-wide patterns of differentiation between the founding resistant and susceptible populations were described using the variants identified by whole genome sequencing of the ten field resistant and ten laboratory susceptible F0 individuals founding the family crosses. A principal components analysis was performed with Plink (v. 1.90b; Chang et al., 2015).

Divergence was measured by Weir and Cockerham’s windowed FST across 40kb windows with a 10kb step using VCFtools (v. 0.1.17; Danecek et al., 2011). A genome wide zFST significance threshold of 6 was used (Rubin et al., 2010). Downstream analysis of this data and all other data presented here were performed in R (v.4.2.1; (R Development Core Team, 2023) with tidyverse R package (v.2.0.0; Wickham et al., 2019), and visualizations were made with the R package ggplot2 (v.3.4.1; Wickham, 2016)

### Quantitative trait mapping for Cry resistance

Cry resistance genomic architecture was characterized by Bayesian sparse linear mixed models (BSLMM) in gemma (v. 0.98.4; Zhou et al., 2013). For each treatment the model was run five separate times, each time with a total of 5 million sampling steps with the first 500 thousand discarded as burn-in. Hyperparameter estimates for each treatment were averaged across all five runs. The association between SNPs across the genome and weight phenotype from the resistance and control growth assays were detected with linear mixed models (LMM) also in gemma (v. 0.98.4; Zhou & Stephens, 2012). Both the BSLMM and LMM account for relatedness among the five replicated families for each treatment.

### Differential gene expression analysis

In 2022, Cry1A.105 + Cry2Ab2 expressing corn ears and ears from the non-Bt expressing near isoline were collected from the University of Maryland CMREC farm at Beltsville. Ears that were infested with *H. zea* were identified by the loosened silk and brought to the laboratory. In the lab, each corn ear was open and any 5th instar larvae present were identified and removed from the ear; caterpillars at other developmental stages were not included in the experiment. Immediately following removal, larvae were chilled and immobilized in a 70% solution of ice cold RNAlater (Invitrogen, Waltham, MA) and PBS. The head capsule on each larva was removed and stored for later measurement to confirm instar. All larvae were later confirmed as 5th instar by head measurements between 1.60 to 2.70 mm (Bilbo et al., 2019). The midgut was cut away from the crop and intestine, and the malpighian tubules were removed.

Corn was cleared from midgut using forceps and a rinse of 70% RNAlater. Midguts were flash frozen and stored at -80℃. The dissection process was repeated with individuals from the laboratory susceptible colony, which had been reared on a standard laboratory diet not containing any corn tissue or Bt toxins.

Midguts of 5th instars from Bt expressing corn, non-Bt corn and the susceptible laboratory colony were organized into eight randomized pools per treatment with four samples in each pool (n = 32 per treatment). RNA was extracted from each pool using am RNeasy® Mini Kit (Qiagen, Hilden, Germany) according to the manufacturer protocol except for the initial pooling of samples. Poly-A selection, library preparation, and sequencing were completed by the University of Maryland Baltimore Genomics Core Facility. All RNA libraries had a RIN value between 8.7 and 10, with 22/24 having RIN of 10, indicating very high quality. Libraries were 100 bp paired end sequenced on an Illumina NovaSeq 6000.

Raw reads were quality controlled with Trimmomatic (v. 0.39; Bolger et al., 2014) to remove Illumina adapters, trim reads where 4 bp sliding window quality score fell below 20, and drop any reads shorter than 36 bp after trimming. The paired quality controlled reads were aligned to a chromosome scale *H. zea* assembly (Benowitz et al., 2022; GCA_022343045.1) with HISAT2 (v. 2.2.1; Kim et al., 2019) using default parameters. Gene annotations were lifted from the older reference genome Hzea_1.0 (Pearce et al., 2017; GCA_002150865.1) to the new chromosome scale assembly using Liftoff (v. 1.6.1; Shumate & Salzberg, 2021). Gene expression counts were generated with HTSeq-count (v. 0.13.5; Anders et al., 2015). Expression patterns across all 24 pools were compared through differential expression analysis in DESeq2 (v. 1.38.3; Love et al., 2014). Two-way differential expression analysis was conducted between resistant field individuals collected from Bt expressing corn and susceptible lab colony samples to identify expression changes associated with resistance. An additional two-way differential expression analysis between the field collected individuals from Bt expressing corn and non expressing corn was conducted to identify Cry exposure inducible gene expression changes. Statistical significance was indicated by a false discovery rate adjusted p-value of 0.01.

### Structural variant detection

Sequencing coverage depth signals were extracted from aligned .bam files from the WGS of the cross founders, the ddRAD sequencing of F2, and two publicly available data sets. The genome wide coverage depth and the average coverage depth for the trypsin cluster were calculated with Qualimap (v. 2.2.1; Okonechnikov et al., 2016), while windowed coverage depth calculations were from samtools depth (v. 1.10; Danecek et al., 2021) results. Calculations of mean coverage of the amplified region for these analyses included all reads aligning between 5.22 - 5.37 Mb on Chr 9. Significant differences in coverage depth across samples were determined by pairwise t-tests with a bonferroni correction to p. The presence of the structural variant in 2002, 2012, and 2017 was assessed using the publicly available whole genome sequencing data from Taylor et al. (2021) (PRJNA751583). The presence of the structural variant in samples collected from Cry expressing and non-expressing corn was assessed using sequencing data from the Pezzini et al. (2024) (PRJNA1055981).

The structural variant was validated with digital droplet PCR (ddPCR) for *tryp77*, one of seven trypsins in the putative duplication. ATP dependent DNA helicase was used as a control gene, as it is single copy in Lepidoptera in OrthoDB (Zdobnov et al., 2021). Twenty-five ng of DNA per sample was mixed with HindIII and analyzed in multiplexed assay for *tryp77* and ATP dependent DNA helicase at MOgene (St. Louis, MO). The genes were targeted with the primers shown in **(Table S10)** and the following amplification conditions: activation at 95℃ for 10 minutes, followed by 40 cycles of denaturing at 94℃ for 30 seconds, annealing and extension at 58℃ for 1 minute, finally the enzyme was deactivated at 98℃ for 10 minutes and held at 4℃. ddPCR was performed with QX200 Automated Droplet Generator and Reader and analyzed using the QX Manager 1.2 Standard Edition software (Bio-Rad, Hercules CA). All samples were run with a synthesized gBlock positive control and a no template negative control. Copy number for samples was calculated as in Karlin-Neumann et al. (2012) with the following formula:

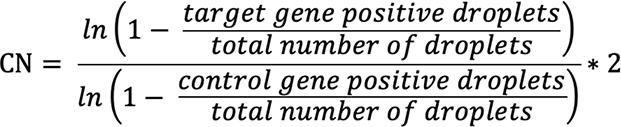

### Field resistant colony collection and rearing

Cry resistant *H. zea* late instar caterpillars were collected from Cry1Ab expressing sweet corn (‘BC0805’) ears grown at the Central Maryland Research and Education Center in Beltsville, Maryland. Approximately 450 caterpillars were collected during each of two collection dates (August 18 and September 7, 2023). Caterpillars were reared on *H. zea* diet (Southland Products Inc., Arkansas, USA) until adulthood and bucket mated in a growth chamber set at 25°C with 70% relative humidity and 16:8 light:dark cycle. Eggs from bucket matings were placed onto diet and reared until the second instar under the same conditions.

Occasionally, larvae were held at 4°C once they reached the appropriate size to ensure sufficient numbers at the same developmental stage for assays.

### Trypsin inhibition assay

To determine whether trypsin activity impacts Cry toxicity in *H. zea*, we compared 7-day larval weight after exposure to corn leaf tissue incorporated diet. Larvae were exposed to one of 12 treatments: 1) 1X TLCK trypsin inhibitor and Cry1Ab expressing leaf tissue, 2) 2X TLCK trypsin inhibitor and Cry1Ab expressing leaf tissue, 3) 5X TLCK trypsin inhibitor and Cry1Ab expressing leaf tissue, 4) 10X TLCK trypsin inhibitor and Cry1Ab expressing leaf tissue, 5) 1X TLCK trypsin inhibitor and non-expressing leaf tissue, 6) 2X TLCK trypsin inhibitor and non-expressing leaf tissue, 7) 5X TLCK trypsin inhibitor and non-expressing leaf tissue, 8) 10X TLCK trypsin inhibitor and non-expressing leaf tissue, 9) buffer and Cry1Ab expressing leaf tissue, 10) buffer and non-expressing leaf tissue, 11) Cry1Ab expressing leaf tissue alone, 12) non-expressing leaf tissue alone. These treatments enabled us to separately determine the growth effect of Cry expressing tissue, the buffer, and the inhibitor at different concentrations. Second instars from the laboratory Cry susceptible *H. zea* population from Benzon research and field-collected Cry resistant *H. zea* were reared on these treatments for seven days before they were individually weighed to assess growth. In each assay, 8 - 16 individuals were exposed to each treatment, due to availability of appropriately sized larvae. Final sample sizes were ≥ 70 in each treatment, and are reported in Table S8.

Based upon Ma et al. (2013) and preliminary trials, we selected N-α-tosyl-ւ-lysine chloromethyl ketone hydrochloride (TLCK, Sigma-Aldrich®, St. Louis, MO) as our trypsin inhibitor. To make the inhibitor solutions, 350 mg of TLCK was dissolved in 10 mL of phosphate buffer solution (0.1M, pH 5.8, bioWORLD, Dublin, OH) to make a stock solution of 35 mg/mL (10X). This solution was further diluted to 3.5 mg/mL (1X), 7 mg/mL (2X), and 17.5 mg/mL (5X) as needed. The inhibitor solutions were freshly made for each assay date. A 10 mL aliquot of phosphate buffer was used for the buffer control treatment. The bioassays were prepared as in Dively (2016), Taylor et al. (2021), and as described above, except, in the inhibitor and buffer treatments 1.6 mL of the TLCK solution or phosphate buffer were mixed into 25 mL of experimental diet, resulting in the following final TLCK concentrations: 1X: 224.5 μg of TLCK per mL diet, 2X: 449 μg of TLCK per mL diet, 5X: 1.1 mg of TLCK per mL diet, 10X: 2.2 mg of TLCK per mL diet.

First we compared the two control groups (incorporated leaf tissue only and incorporated leaf tissue with buffer) using t tests with an α of 0.01 to identify any potential effects of the buffer alone. Then, we tested for a synergistic effect of trypsin inhibition and Cry exposure on growth in *H. zea* with a general linear model comparison approach. A gamma distribution was used for the weight response variable. The fit of the models was compared with a likelihood ratio chi-squared test using an α of 0.01. We tested for combined effects by comparing a full model with Cry treatment and trypsin inhibition main effects and an interaction between the two, to a model with only the two main effects. Bonferroni corrected post hoc contrasts were calculated with the R package emmeans (v. 1.8.6; Lenth et al., 2023). We also calculated a growth ratio at each TLCK dose as:

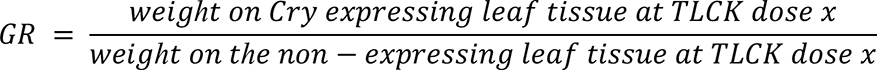

The growth ratio measured the combined effect of TLCK and Cry toxins, and would be expected to stay consistent across doses if the effects of TLCK and Cry toxins on growth suppression are not linked.

## Acknowledgements

We thank Juan Luis Jurat-Fuentes, and David Heckel for suggestions that improved our manuscript. We thank Galen Dively for access to experimental plots for field collections. We thank Katherine Bell, Thea Bliss, Ben Burgunder, Lasair ní Chochlain, Maria Cramer, Kyree Day, Dominique Desmarattes, Shea Ill, Emma Kohanski, Ava Lamberty, Scott McCluen, Declan McHugh, Oliva Moy, Hiral Patel, Daniella Anconeira Sayco, Rachel Sanford, Torsten Schöneberg, Robert Starkenburg, Olivia Rosen, and Joshua Yeroshefsky for assistance with H. zea collections, crosses, rearing, bioassays, and/or DNA isolation. This work was funded by the US Department of Agriculture, National Institute of Food and Agriculture Biotechnology Risk Assessment Grants 2019-33522-29992 and 2022-33522-37744.

## Notes

### Competing Interest Statement

The authors have declared no competing interest.

### Summary of Updates

A new analysis of publicly available WGS of field samples has been completed. Other minor new analyses added. The manuscript has been updated to reflect these changes and edited for clarity throughout.

