## Supporting Information for "Polygenic basis of Bt Cry resistance evolution in wild *Helicoverpa zea*"

**Supporting Information - Taylor et al. 2024**

### Supplementary Results

*Cry resistance phenotypes*

A stepwise model reduction and comparison approach showed that treatment (Cry1Ab, Cry1A.105 + Cry2Ab2, or their respective controls), population ($F_0$ resistant, $F_0$ susceptible, or $F_2$), and the interaction between treatment and population all significantly improved the fit of the model. Pairwise post hoc contrasts of the estimated marginal means revealed significant differences (bonferroni adjusted $p < 0.01$) for all comparisons between populations in each treatment, meaning that $F_0$ resistant, $F_0$ susceptible, and $F_2$ intercross offspring growth differences were significant in both Cry expressing and control treatments (**Table S1, Table S2)**. In all treatments, the resistant founding parents grew the most, while the $F_2$ offspring had intermediate growth and the susceptible population grew the least (**Figure 1**). Post hoc tests also revealed that all comparisons between treatments for each population were significantly different (bonferroni adjusted $p < 0.01$), except for the comparisons between the two control growth treatments for both founding ($F_0$) populations (**Table S1, Table S2)**. For all populations, the Cry1A.105 + Cry2Ab2 treatment restricted growth more than the Cry1Ab treatment, and individuals grew largest on the control growth assays without toxin exposure (**Figure 1**).

Resistant individuals that founded the crosses ranged in weight from 81.2 – 112.7 mg on the Cry1Ab containing leaf treatment, and 58.9 – 92.9 mg on the Cry1A.105 + Cry2Ab2 expressing leaf treatment. The founding resistant parent of each cross was selected to have a growth phenotype that was above the first quartile. The susceptible lab colony growth phenotypes were on average 16.7 mg in the Cry1Ab expressing treatment and 4.8 mg in the Cry1A.105 + Cry2Ab2 expressing treatment. The intermediate $F_2$ phenotypes, the rarity of $F_2$ offspring recovering the most extreme founding population phenotypes, and the somewhat normal appearing phenotypic distributions, suggested that both Cry resistance and general growth in the laboratory assays were quantitative genetic traits (**Figure S1**). We also observed variation in both growth and Cry resistance phenotypes among families, suggesting that mapping families and the resistant field parent who founded them varied in which resistance and growth associated traits they possessed (**Figure S1**).

*Genome wide divergence between resistant and susceptible H. zea*

Genome wide divergence was high between the resistant and susceptible cross founding populations. Strong signals of divergence were expected due to the long-term genetic isolation between the field resistant and susceptible laboratory populations used in our study, and suggest the necessity of laboratory cross experiments for the detection of genomic regions involved in resistance. In a principal components analysis of genome wide SNPs, the first principal component explained 38.8% of variance in the dataset and clearly separated resistant field and laboratory susceptible cross parents. Variation between field resistant parents could be attributed to principal component 2, which explained 11.4% of variance (**Figure S2)**. We also measured genome wide divergence using Weir and Cockerham's weighted $F_{ST}$ calculated genome wide in

40kb windows with a 10kb step. Divergence was high between resistant and susceptible populations with a mean windowed $F_{ST}$ of 0.40. Variance in $F_{ST}$ across the genome was wide with a range of window means from -0.05 to 0.89. Though the populations show strong signals of genome-wide divergence, no windows fell above a genome-wide $zF_{ST}$ threshold of 6 (Rubin et al., 2010) (**Figure S3**). We also observed a large number of SNPs fixed between resistant and susceptible populations (**Figure S3**). In both this analysis and Taylor et al. (2021) the windows with highest weighted $F_{ST}$ genome wide were within 10 Kb of 4 Mb on Chr 13; the peak $F_{ST}$ window for the resistant and susceptible parent comparison was from 4.002 - 4.042 Mb, and for time comparison it was from 3.958 - 3.998 Mb. Additionally, an overlapping peak is found at 5.75 Mb on Chr 9, a chromosome we previously linked to Cry resistance phenotypes (**Figure S3**) (Taylor et al. 2021). These consistent patterns suggest that the susceptible population has maintained historical susceptible genotypes.

*Evidence for roles of other major candidate genes*

We searched for other major Cry resistance candidates in our large QTL regions. As in Taylor et al. 2021, we found that the top candidate genes discussed in the literature did not fall in our detected QTL even with an increased sample size and replicated mapping families. These candidates included ABCA2, ABCC2, ABCG1, CAD2, CAD86C, MAP4K4, TSPAN1 (Fabrick et al., 2022; Fritz et al., 2020; Gahan et al., 2010; Gill et al., 1995; Guo, Kang, Chen, et al., 2015; Guo, Kang, Zhu, et al., 2015; Tay et al., 2015; Z. Zhang et al., 2017). The candidate genes ALP (Jurat-Fuentes & Adang, 2004) and CALP (Gahan et al., 2001) were found on chromosomes associated with resistance, but were not in the regions most strongly associated with phenotype. A cluster of candidate aminopeptidases (Banks et al., 2001) were found on Chr 9, but were also not near the QTL peak. We previously showed that variant frequencies within these candidate genes did not change significantly while Cry resistance was evolving (Taylor et al., 2021). Based upon our prior results and the lack of strong linkage with phenotype detected here, these candidate genes were unlikely to have a major role in Cry resistance evolution in *H. zea*.

**Supplementary Tables**

**Table S1**. Cry resistance and growth phenotypes in a leaf tissue incorporation lab assay measured as weight after seven days of growth on toxin containing and control diets.

| Population | Treatment | n | Weight mean (mg) | Weight sd |
|---|---|---|---|---|
| $F_0$ Susceptible | Cry1Ab | 187 | 16.7 | 13.4 |
| $F_0$ Resistant | Cry1Ab | 302 | 95.6 | 39.0 |
| $F_2$ | Cry1Ab | 544 | 54.8 | 35.4 |
| $F_0$ Susceptible | Cry1A.105+Cry2Ab2 | 185 | 4.8 | 4.4 |
| $F_0$ Resistant | Cry1A.105+Cry2Ab2 | 299 | 72.8 | 34.9 |
| $F_2$ | Cry1A.105+Cry2Ab2 | 562 | 21.5 | 19.1 |
| $F_0$ Susceptible | Control for Cry1Ab | 90 | 69.7 | 51.2 |
| $F_0$ Resistant | Control for Cry1Ab | 151 | 129.8 | 43.9 |
| $F_2$ | Control for Cry1Ab | 543 | 106.8 | 62.8 |
| $F_0$ Susceptible | Control for Cry1A.105+Cry2Ab2 | 87 | 62.8 | 49.0 |
| $F_0$ Resistant | Control for Cry1A.105+Cry2Ab2 | 153 | 131.5 | 45.6 |
| $F_2$ | Control for Cry1A.105+Cry2Ab2 | 575 | 87.4 | 64.8 |

**Table S2**. Pairwise post hoc comparison of treatment group effects in phenotypic assays of Cry resistance and growth phenotypes. At the top of the table are comparisons between populations (resistant, susceptible, and $F_2$) within a single treatment (Cry1Ab, control for Cry1Ab, Cry1A.105 + Cry2Ab2, and control for Cry1A.105 + Cry2Ab2). Below are the comparisons with a single population but between treatments.

| Group A | Group B | Estimate | DF | p |
|---|---|---|---|---|
| Cry1Ab | | | | |
| $F_0$ Resistant | $F_2$ | -7.78 | 3666 | <.0001 |
| $F_0$ Resistant | $F_0$ Susceptible | -49.3 | 3666 | <.0001 |
| $F_2$ | $F_0$ Susceptible | -41.52 | 3666 | <.0001 |
| Control for Cry1Ab | | | | |
| $F_0$ Resistant | $F_2$ | -1.66 | 3666 | 0.0032 |
| $F_0$ Resistant | $F_0$ Susceptible | -6.64 | 3666 | <.0001 |
| $F_2$ | $F_0$ Susceptible | -4.98 | 3666 | <.0001 |
| Cry1A.105 + Cry2Ab2 | | | | |
| $F_0$ Resistant | $F_2$ | -32.82 | 3666 | <.0001 |
| $F_0$ Resistant | $F_0$ Susceptible | -192.7 | 3666 | <.0001 |
| $F_2$ | $F_0$ Susceptible | -159.88 | 3666 | <.0001 |
| Control B | | | | |
| $F_0$ Resistant | $F_2$ | -3.83 | 3666 | <.0001 |
| $F_0$ Resistant | $F_0$ Susceptible | -8.32 | 3666 | <.0001 |
| $F_2$ | $F_0$ Susceptible | -4.49 | 3666 | 0.0006 |
| $F_0$ Resistant Founding Population | | | | |
| Control for Cry1A.105+Cry2Ab2 | Cry1A.105 + Cry2Ab2 | -6.12 | 3666 | <.0001 |
| Control for Cry1A.105+Cry2Ab2 | Control for Cry1Ab | -0.1 | 3666 | 1 |
| Control for Cry1A.105+Cry2Ab2 | Cry1Ab | -2.86 | 3666 | <.0001 |
| Cry1A.105 + Cry2Ab2 | Control for Cry1Ab | 6.03 | 3666 | <.0001 |
| Cry1A.105 + Cry2Ab2 | Cry1Ab | 3.27 | 3666 | <.0001 |
| Control for Cry1Ab | Cry1Ab | -2.76 | 3666 | <.0001 |
| $F_2$ offspring | | | | |
| Control for Cry1A.105+Cry2Ab2 | Cry1A.105 + Cry2Ab2 | -35.11 | 3666 | <.0001 |
| Control for Cry1A.105+Cry2Ab2 | Control for Cry1Ab | 2.08 | 3666 | <.0001 |
| Control for Cry1A.105+Cry2Ab2 | Cry1Ab | -6.81 | 3666 | <.0001 |
| Cry1A.105 + Cry2Ab2 | Control for Cry1Ab | 37.19 | 3666 | <.0001 |
| Cry1A.105 + Cry2Ab2 | Cry1Ab | 28.31 | 3666 | <.0001 |
| Control for Cry1Ab | Cry1Ab | -8.88 | 3666 | <.0001 |
| $F_0$ Susceptible Founding Population | | | | |

| | | | | |
|---|---|---|---|---|
| Control for Cry1A.105+Cry2Ab2 | Cry1A.105 + Cry2Ab2 | -190.5 | 3666 | <.0001 |
| Control for Cry1A.105+Cry2Ab2 | Control for Cry1Ab | 1.58 | 3666 | 1 |
| Control for Cry1A.105+Cry2Ab2 | Cry1Ab | -43.84 | 3666 | <.0001 |
| Cry1A.105 + Cry2Ab2 | Control for Cry1Ab | 192.09 | 3666 | <.0001 |
| Cry1A.105 + Cry2Ab2 | Cry1Ab | 146.67 | 3666 | <.0001 |
| Control for Cry1Ab | Cry1Ab | -45.42 | 3666 | <.0001 |

**Table S3** Genes annotated on Chromosome 9 between 5 and 6 Mb.

| Start | Stop | +/- | Gene ID | Annotation | Code |
|---|---|---|---|---|---|
| 5018492 | 5018875 | - | HzOG216674 | BMORI:regulating synaptic membrane exocytosis protein 2-like | |
| 5036816 | 5068300 | - | HzOG216675 | BMORI:cytochrome b5-related protein-like | |
| 5047052 | 5049569 | - | HzOG216676 | BMORI:cytochrome b5-related protein-like | |
| 5106488 | 5111179 | - | HzOG216677 | BMORI:cytochrome b5-related protein-like | |
| 5113361 | 5115476 | + | HzOG216678 | DMELA:Q9VJI1 CG17928 | |
| 5133223 | 5134095 | + | HzOG216680 | BMORI:ommochrome-binding protein-like | |
| 5200320 | 5207810 | - | HzOG216681 | BMORI:myrosinase 1-like | Myr1A |
| 5212768 | 5216866 | - | HzOG216682 | BMORI:myrosinase 1-like | Myr1B |
| 5231258 | 5253118 | + | HzOG216683 | BMORI:disks large homolog 4-like | Dlh |
| 5273366 | 5281941 | - | HzOG200427 | HzeaTryp073 | Tryp73 |
| 5289846 | 5293176 | + | HzOG200430 | HzeaTryp076 | Tryp76 |
| 5296095 | 5299655 | + | HzOG200429 | HzeaTryp075 | Tryp75 |
| 5306427 | 5309805 | + | HzOG200431 | HzeaTryp077 | Tryp77 |
| 5313249 | 5316522 | + | HzOG200432 | HzeaTryp078 | Tryp78 |
| 5318668 | 5318763 | + | HzOG202341 | BMORI:neuropeptide receptor A35 | Npr |
| 5330236 | 5340510 | + | HzOG200433 | HzeaTryp079 | Tryp79 |
| 5336767 | 5340510 | + | HzOG200434 | HzeaTryp080 | Tryp80 |
| 5444566 | 5445458 | + | HzOG216686 | BMORI:uncharacterized protein LOC101738586 isoform X2 | |
| 5452851 | 5455682 | - | HzOG216687 | DMELA:Q9VC31 RabX4 BMORI:ras-related protein Rab-8A-like | |
| 5493784 | 5641495 | - | HzOG216688 | BMORI:ras-related and estrogen-regulated growth inhibitor-like protein-like | |
| 5502346 | 5505035 | - | HzOG216690 | BMORI:tubulin glycylase 3A-like | |
| 5563384 | 5571942 | - | HzOG216692 | BMORI:ras-related and estrogen-regulated growth inhibitor-like protein-like | |
| 5582602 | 5583984 | - | HzOG216693 | DMELA:Q9V3R6 tRNA-specific adenosine deaminase 1 | |
| 5608838 | 5609053 | + | HzOG216694 | DMELA:Q4QPR8 CG14483 | |
| 5609339 | 5611366 | - | HzOG216695 | HSAPI:Q8NBP5 Major facilitator superfamily domain-containing protein 9 | |
| 5616334 | 5617499 | + | HzOG216696 | DMELA:Q7PL91 CG40002 | |
| 5630362 | 5636972 | + | HzOG216697 | BMORI:SH3 domain-containing kinase-binding protein 1-like | |
| 5640630 | 5644033 | - | HzOG216698 | HSAPI:A6NG64 Leucine-rich repeat-containing protein C10orf11 | |
| 5647703 | 5653476 | - | HzOG216699 | BMORI:retinal dehydrogenase 1-like | |
| 5657464 | 5662801 | - | HzOG216700 | BMORI:retinal dehydrogenase 1-like | |
| 5664173 | 5674759 | - | HzOG216701 | retinal dehydrogenase 1-like | |
| 5670410 | 5674759 | - | HzOG216702 | retinal dehydrogenase 1-like | |

| | | | | |
|---|---|---|---|---|
| 5697431 | 5702619 | + | HzOG216703 | DMELA:P25931 Neuropeptide Y receptor |
| 5745541 | 5760783 | - | HzOG206139 | BMORI:neuropeptide receptor A19 |
| 5834992 | 5836132 | - | HzOG206140 | BMORI:uncharacterized protein LOC101741733 |
| 5839370 | 5885612 | + | HzOG206141 | BMORI:uncharacterized protein LOC101744381 |
| 5842612 | 5846718 | - | HzOG206142 | BMORI:uncharacterized protein LOC101741593 |
| 5847697 | 5848764 | + | HzOG206143 | DMELA:Q95RV5 Ribulose-phosphate 3-epimerase BMORI:ribulose-phosphate 3-epimerase-like |
| 5848900 | 5850280 | - | HzOG206144 | DMELA:A1Z830 CG12128, isoform A |
| 5863186 | 5870131 | + | HzOG206145 | DMELA:Q95RT1 LD12501p |
| 5873037 | 5892506 | + | HzOG206146 | Rel [FBgn0014018] |
| 5894662 | 5897463 | + | HzOG206147 | origin recognition complex subunit 2-like |
| 5899282 | 5903078 | - | HzOG206148 | DMELA:Q9V895 Acidic leucine-rich nuclear phosphoprotein 32 family member |
| 5920200 | 5921378 | + | HzOG206149 | BMORI:alpha-1,6-mannosyl-glycoprotein 2-beta-N-acetylglucosaminyltransferase-like |

**Table S4** Genes annotated on Chromosome 30 between 2.4 - 3.4 Mb.

| Start | Stop | +/- | Gene ID | Annotation |
|---|---|---|---|---|
| 2409534 | 2411162 | + | HzOG215027 | BMORI:lysosome membrane protein 2-like |
| 2422680 | 2426798 | + | HzOG215447 | ScR-B13 |
| 2422680 | 2426798 | + | HzOG215876 | ScR-B4 |
| 2429256 | 2440911 | - | HzOG215875 | BMORI:putative fatty acyl-CoA reductase CG5065-like |
| 2431403 | 2441394 | - | HzOG215448 | BMORI:putative fatty acyl-CoA reductase CG5065-like |
| 2446567 | 2457167 | - | HzOG215449 | BMORI:putative fatty acyl-CoA reductase CG5065-like |
| 2486261 | 2491502 | - | HzOG215703 | DMELA:Q9VPH1 Toll-9 |
| 2487234 | 2500480 | - | HzOG215377 | Toll9 |
| 2501683 | 2509177 | + | HzOG215378 | DMELA:A8JUM9 CG15373, isoform B |
| 2517814 | 2522906 | - | HzOG215379 | DMELA:Q7KK29 BcDNA.LD27979 |
| 2528138 | 2532944 | - | HzOG214905 | BMORI:GATA zinc finger domain-containing protein 14-like |
| 2539815 | 2542223 | - | HzOG200163 | HzeaCCE019b |
| 2553650 | 2556859 | + | HzOG200162 | HzeaCCE019a |
| 2560776 | 2568608 | - | HzOG200161 | HzeaCCE014a |
| 2598623 | 2603529 | + | HzOG215396 | BMORI:uncharacterized protein LOC101739044 |
| 2631989 | 2634956 | - | HzOG215307 | BMORI:uncharacterized protein LOC101745937 |
| 2645025 | 2668686 | + | HzOG200345 | HzeaABCC11 |
| 2674016 | 2681372 | - | HzOG214993 | BMORI:LOW QUALITY PROTEIN: twitchin-like |
| 2696103 | 2697701 | - | HzOG214995 | acetyl-CoA carboxylase-like |
| 2702471 | 2705627 | - | HzOG216185 | acetyl-CoA carboxylase-like |
| 2708232 | 2714004 | + | HzOG200982 | HzeaOR20 |
| 2748213 | 2748476 | - | HzOG215302 | BMORI:acetyl-CoA carboxylase-like |
| 2796129 | 2808941 | - | HzOG210885 | BMORI:P protein-like |
| 2825587 | 2847206 | - | HzOG210883 | BMORI:P protein-like |
| 2854428 | 2872148 | - | HzOG210882 | BMORI:IQ and AAA domain-containing protein 1-like |
| 2878310 | 2882055 | - | HzOG210881 | BMORI:TBC1 domain family member 1-like |
| 2896367 | 2902406 | - | HzOG210880 | DMELA:E1JJ52 FI17814p1 |

| | | | | |
|---|---|---|---|---|
| 2935517 | 2938166 | - | HzOG210878 | BMORI:transmembrane protein 214-like isoform X2 |
| 2961017 | 2979003 | + | HzOG210877 | DMELA:Q8I930 GH14147p |
| 3178539 | 3231243 | - | HzOG215010 | BMORI:RNA-binding protein 26-like |
| 3178947 | 3182843 | + | HzOG216180 | BMORI:uncharacterized protein LOC101741223 isoform X1 |
| 3197929 | 3201348 | - | HzOG212889 | BMORI:RNA-binding protein 26-like |
| 3271854 | 3286917 | + | HzOG200933 | HzeaGR204 |
| 3280084 | 3281271 | - | HzOG212873 | BMORI:uncharacterized protein LOC101741081 |
| 3283456 | 3284649 | - | HzOG212872 | BMORI:uncharacterized protein LOC101740334 |
| 3294683 | 3306849 | - | HzOG212871 | MAP kinase-activated protein kinase 2-like |
| 3318996 | 3319072 | - | HzOG206936 | BMORI:fukutin-related protein-like |
| 3382826 | 3387263 | - | HzOG200395 | HzeaTryp129 |

**Table S5.** Top 50 differentially expressed genes in resistant field samples relative to susceptible laboratory controls ranked from smallest to largest adjusted p-value (all < 0.001). Positive values indicate up regulation in the resistant population while negative values indicate down regulation in the resistant population.

| Chrom | Gene ID | Start | End | Base Mean Expression | Log2 Fold Change | Gene Annotation |
|---|---|---|---|---|---|---|
| Hz_05 | HzOG200462 | 11357047 | 11358145 | 17450 | 9.6 | HzeaTryp119 |
| Hz_23 | HzOG211681 | 7409556 | 7409813 | 2542 | 5.2 | No annotation - blast hit to SFRUG: histidine-rich glycoprotein-like |
| Hz_22 | HzOG201840 | 8160815 | 8163190 | 447 | 3 | HSAPI:Q9Y315 Putative deoxyribose-phosphate aldolase |
| Hz_15 | HzOG200202 | 9489071 | 9491580 | 92728 | 5.3 | HzeaCCE001g |
| Hz_09 | HzOG200431 | 5306427 | 5309805 | 108320 | 3.8 | HzeaTryp077 |
| Hz_18 | HzOG203325 | 8849649 | 8849993 | 129 | -6.7 | BMORI:uncharacterized protein LOC101744851 |
| Hz_23 | HzOG200149 | 1276950 | 1278197 | 1180 | -7.8 | HzeaCCE016i |
| Hz_15 | HzOG206785 | 3070527 | 3072844 | 25481 | 6.9 | BMORI:hatching enzyme-like II precursor |
| Hz_05 | HzOG200456 | 11384357 | 11385400 | 6573 | 7 | HzeaTryp113 |
| Hz_15 | HzOG215780 | 4754409 | 4763551 | 94 | -5.2 | BMORI:uncharacterized protein LOC101735407 |
| Hz_09 | HzOG200432 | 5313249 | 5316522 | 97882 | 3.9 | HzeaTryp078 |
| Hz_09 | HzOG200434 | 5336767 | 5340510 | 49473 | 4.3 | HzeaTryp080 |
| Hz_03 | HzOG203362 | 11306004 | 11307528 | 228 | 3.5 | DMELA:Q7KRY6 Nucleosomal histone kinase 1 |
| Hz_05 | HzOG200457 | 11378257 | 11379712 | 374 | 9.6 | HzeaTryp114 |
| Hz_09 | HzOG200427 | 5273366 | 5281941 | 109480 | 3 | HzeaTryp073 |
| Hz_07 | HzOG215059 | 1524496 | 1539112 | 154 | -9.3 | DMELA:Q9VC92 CG6432 |
| Hz_20 | HzOG211283 | 5352289 | 5353116 | 31955 | 5.3 | BMORI:probable salivary secreted peptide-like |
| Hz_09 | HzOG200430 | 5289846 | 5293176 | 25248 | 3.8 | HzeaTryp076 |

| | | | | | | |
|---|---|---|---|---|---|---|
| Hz_10 | HzOG212164 | 4276503 | 4277474 | 45 | 3.4 | BMORI:probable ATP-dependent RNA helicase ddx17-like |
| Hz_05 | HzOG205041 | 14634337 | 14635410 | 145 | 5.7 | BMORI:uncharacterized protein LOC101736412 |
| Hz_09 | HzOG216682 | 5212768 | 5216866 | 126 | 7.6 | BMORI:myrosinase 1-like |
| Hz_21 | HzOG200515 | 10494801 | 10501265 | 60514 | 4.2 | HzeaChym018 |
| Hz_31 | HzOG205106 | 1289550 | 1291313 | 24232 | -2.9 | BMORI:cadherin-87A-like, partial |
| Hz_30 | HzOG205921 | 6504686 | 6509827 | 9035 | 2.9 | BMORI:leukocyte surface antigen CD53-like isoform X2 |
| Hz_09 | HzOG200429 | 5296095 | 5299655 | 55 | 5.5 | HzeaTryp075 |
| Hz_20 | HzOG216440 | 3852789 | 3853472 | 39 | 10.4 | BMORI:uncharacterized protein LOC101735858 |
| Hz_14 | HzOG203230 | 10815828 | 10817428 | 53 | -8.3 | BMORI:uncharacterized protein LOC101744346 |
| Hz_30 | HzOG200163 | 2539815 | 2542223 | 168 | 2.1 | HzeaCCE019b |
| Hz_28 | HzOG205157 | 7369695 | 7373135 | 168 | -10 | BMORI:coiled-coil domain-containing protein 19, mitochondrial-like |
| Hz_20 | HzOG213201 | 5903806 | 5910406 | 55 | 3.7 | BMORI:uncharacterized protein LOC101745265 |
| Hz_16 | HzOG210085 | 10926011 | 10928082 | 43 | -8.3 | BMORI:alpha-tocopherol transfer protein-like |
| Hz_11 | HzOG210674 | 1081425 | 1098906 | 24765 | 2.3 | BMORI:gelsolin, cytoplasmic-like |
| Hz_05 | HzOG200450 | 11413496 | 11414573 | 169 | 3.7 | HzeaTryp107 |
| Hz_19 | HzOG200675 | 6751266 | 6752714 | 889 | 5.2 | HzeaCSP12 |
| Hz_04 | HzOG203617 | 546035 | 549917 | 7422 | 5.7 | BMORI:maltase 1-like |
| Hz_15 | HzOG213605 | 13079027 | 13081145 | 8058 | 6.3 | BMORI:probable salivary secreted peptide-like |
| Hz_07 | HzOG204020 | 5522296 | 5538830 | 40075 | 4.8 | BMORI:uncharacterized protein LOC101737697 |
| Hz_03 | HzOG213452 | 9870099 | 9872491 | 303 | 4.5 | BMORI:glyoxylate reductase/hydroxypyruvate reductase-like |

| | | | | | | |
|---|---|---|---|---|---|---|
| Hz_25 | HzOG204807 | 9339936 | 9342133 | 77 | 6.3 | BMORI:probable E3 ubiquitin-protein ligase HERC2-like |
| Hz_30 | HzOG205951 | 5263612 | 5270482 | 26272 | 0.8 | BMORI:leucine-rich repeat neuronal protein 3-like isoform X3 |
| Hz_08 | HzOG207928 | 417041 | 420971 | 142 | 4.4 | BMORI:putative inorganic phosphate cotransporter-like |
| Hz_14 | HzOG200005 | 6541086 | 6553353 | 41 | -4.2 | HzeaCYP301B1 |
| Hz_01 | HzOG215482 | 14048149 | 14051195 | 7642 | 1.4 | BMORI:uncharacterized protein LOC101740601 |
| Hz_01 | HzOG201736 | 14897993 | 14898526 | 43 | -8.1 | BMORI:uncharacterized protein LOC101742112 |
| Hz_07 | HzOG200381 | 1317173 | 1337872 | 19301 | 5.5 | HzeaTryp027 |
| Hz_29 | HzOG207844 | 5365418 | 5367481 | 684 | 3.1 | BMORI:peroxisomal membrane protein PMP34-like |
| Hz_05 | HzOG200449 | 11413496 | 11418142 | 1692 | 3.5 | HzeaTryp106 |
| Hz_25 | HzOG200118 | 1925382 | 1927252 | 438 | 6 | HzeaCCE006i |
| Hz_13 | HzOG207750 | 3758456 | 3759224 | 282 | -4.5 | BMORI:1-acylglycerophosphocholine O-acyltransferase 1-like |
| Hz_20 | HzOG210545 | 8848940 | 8849116 | 61 | 5.6 | BMORI:vacuolar protein sorting-associated protein 35-like |

**Table S6.** Significantly differentially expressed genes (adjusted p-value < 0.01) in resistant field samples relative to susceptible laboratory controls that are found within 100 kb of a SNP significantly associated (adjusted p-value < 0.01) with resistance phenotype in the QTL study. SNP association is the treatment(s) in which the nearby resistance associated SNP was identified. Positive values indicate up regulation in the resistant population while negative values indicate down regulation in the resistant population.

| SNP Association | Chr | Gene ID | Start | End | Base Mean | Log2 Fold Change | Gene Annotation |
|---|---|---|---|---|---|---|---|
| Cry1 | Hz_02 | HzOG203798 | 4501521 | 4501747 | 38.80 | 1.161 | BMORI:histidine-rich glycoprotein-like |
| Cry1 | Hz_02 | HzOG200841 | 6875308 | 6879425 | 798.74 | 3.604 | HzeaOBP14 |
| Cry1 | Hz_03 | HzOG214451 | 6278328 | 6308682 | 97.93 | -1.736 | DMELA:Q494G1 CG9864 |
| Cry1 | Hz_03 | HzOG217072 | 7218588 | 7220526 | 943.64 | -1.174 | BMORI:uncharacterized protein LOC101746778 |
| Cry1+2 | Hz_09 | HzOG214291 | 514101 | 515235 | 58.33 | 5.117 | BMORI:LOW QUALITY PROTEIN: diacylglycerol kinase 1-like |
| Cry1+2 | Hz_09 | HzOG214290 | 517186 | 519903 | 7.55 | 4.394 | BMORI:contactin-like |
| Cry1 | Hz_09 | HzOG214262 | 1054642 | 1062456 | 46.84 | 1.122 | BMORI:LOW QUALITY PROTEIN: myotubularin-related protein 5-like |
| Cry1 | Hz_09 | HzOG214257 | 1102040 | 1104152 | 61.46 | 1.567 | Eater2 |
| Cry1 | Hz_09 | HzOG215623 | 1916962 | 1918403 | 23.23 | 4.613 | BMORI:uncharacterized protein LOC101735735 |
| Cry1+2 | Hz_09 | HzOG216612 | 3540933 | 3547901 | 62.92 | -2.249 | DMELA:O18475 FI03732p |
| Cry1 | Hz_09 | HzOG216665 | 4526572 | 4534288 | 127.12 | -2.145 | DMELA:Q7KSA4 CG5191, isoform C |
| Both | Hz_09 | HzOG216682 | 5212768 | 5216866 | 125.60 | 7.623 | BMORI:myrosinase 1-like |
| Both | Hz_09 | HzOG200427 | 5273366 | 5281941 | 109479.50 | 2.971 | HzeaTryp073 |
| Both | Hz_09 | HzOG200430 | 5289846 | 5293176 | 25247.64 | 3.835 | HzeaTryp076 |
| Both | Hz_09 | HzOG200429 | 5296095 | 5299655 | 54.69 | 5.483 | HzeaTryp075 |
| Both | Hz_09 | HzOG200431 | 5306427 | 5309805 | 108319.80 | 3.785 | HzeaTryp077 |
| Both | Hz_09 | HzOG200432 | 5313249 | 5316522 | 97882.06 | 3.878 | HzeaTryp078 |
| Both | Hz_09 | HzOG200433 | 5330236 | 5340510 | 20639.62 | 1.859 | HzeaTryp079 |
| Both | Hz_09 | HzOG200434 | 5336767 | 5340510 | 49473.16 | 4.323 | HzeaTryp080 |
| Both | Hz_09 | HzOG206156 | 6064366 | 6067010 | 75.41 | -1.778 | BMORI:uncharacterized protein LOC101742906 |
| Cry1 | Hz_09 | HzOG206163 | 6190124 | 6192892 | 404.49 | 0.592 | DMELA:Q86P44 SD27779p |

| | | | | | | | |
|---|---|---|---|---|---|---|---|
| Both | Hz_09 | HzOG206206 | 7754662 | 7755451 | 1350.39 | -0.844 | DMELA:P49963 Signal recognition particle 19 kDa protein |
| Both | Hz_09 | HzOG206225 | 8026790 | 8028219 | 704.80 | -1 | HzeaRPL32m |
| Both | Hz_09 | HzOG206282 | 9308409 | 9312863 | 11.50 | 0.034 | BMORI:arylsulfatase B-like |
| Both | Hz_09 | HzOG206301 | 9793491 | 9794774 | 36.67 | -0.792 | HSAPI:P51159 Ras-related protein Rab-27A |
| Both | Hz_09 | HzOG200780 | 10030438 | 10038009 | 3.46 | -4.223 | HzeaOR42 |
| Cry1 | Hz_09 | HzOG205272 | 10058928 | 10060580 | 218.83 | 1.118 | HSAPI:Q9NXA8 NAD-dependent protein deacylase sirtuin-5, mitochondrial |
| Cry1 | Hz_09 | HzOG205140 | 11389070 | 11399696 | 214959.20 | 1.368 | HaAPN1 |
| Cry1 | Hz_09 | HzOG211630 | 11711213 | 11715165 | 68.97 | 1.598 | BMORI:WAP four-disulfide core domain protein 8-like |
| Cry1 | Hz_09 | HzOG206009 | 12316252 | 12320841 | 1683.81 | -1.254 | DMELA:A8QHW8 CG17691, isoform F |
| Cry1 | Hz_09 | HzOG213180 | 12403898 | 12409620 | 51.43 | -1.134 | BMORI:meiosis-specific nuclear structural protein 1-like |
| Cry1 | Hz_09 | HzOG213356 | 12942114 | 12942542 | 10.11 | -3.556 | BMORI:uncharacterized protein LOC101735917 |
| Cry1 | Hz_30 | HzOG204691 | 4080828 | 4082487 | 3.36 | 3.034 | BMORI:uncharacterized protein LOC101736946 |

**Table S7.** All significantly differentially expressed genes in resistant field samples collected from Cry expressing corn relative to field samples collected corn not expressing Cry toxins, ranked from smallest to largest adjusted p-value (all < 0.01). Positive values indicate up regulation in the resistant Cry exposed population while negative values indicate down regulation in the resistant Cry exposed population.

| Chrom | Gene ID | Start | End | Base Mean Expression | Log2 Fold Change | Gene Annotation |
|---|---|---|---|---|---|---|
| Hz_05 | HzOG200461 | 11361437 | 11362474 | 2570 | 5.063 | HzeaTryp118 |
| Hz_15 | HzOG206785 | 3070527 | 3072844 | 25481 | 3.987 | BMORI:hatching enzyme-like II precursor |
| Hz_13 | HzOG215003 | 5560641 | 5568051 | 15 | 0.002 | BMORI:myosin-9-like |
| Hz_12 | HzOG213727 | 2498045 | 2499459 | 13459 | -2.848 | Lysozyme1 |
| Hz_25 | HzOG202191 | 2011243 | 2011986 | 226 | -3.507 | BMORI:uncharacterized protein LOC101743112 |
| Hz_29 | HzOG200554 | 6023113 | 6025984 | 31 | 3.525 | HzeaLipase27 |
| Hz_20 | HzOG200153 | 7870386 | 7872836 | 964 | -3.185 | HzeaCCE006d |
| Hz_16 | HzOG207618 | 8706850 | 8710026 | 716 | -0.101 | BMORI:phosphatidylethanolamine-binding protein homolog F40A3.3-like isoform X3 |
| Hz_05 | HzOG200609 | 5066815 | 5068214 | 30889 | 2.438 | HzeaLipase82 |
| Hz_31 | HzOG205106 | 1289550 | 1291313 | 24232 | -1.497 | BMORI:cadherin-87A-like, partial |
| Hz_14 | HzOG212920 | 5563526 | 5567610 | 1 | -0.002 | BMORI:pickpocket protein 28-like |
| Hz_07 | HzOG215055 | 1596747 | 1612399 | 355 | -0.928 | BMORI:myosuppressin precursor |
| Hz_05 | HzOG200608 | 5059409 | 5061274 | 8326 | 2.15 | HzeaLipase81 |
| Hz_05 | HzOG200448 | 11420529 | 11421636 | 43 | 2.644 | HzeaTryp105 |
| Hz_02 | HzOG202622 | 376269 | 377565 | 511 | -0.746 | BMORI:uncharacterized protein LOC101742051 |
| Hz_15 | HzOG214752 | 12671492 | 12678231 | 222 | 1.726 | BMORI:uncharacterized protein LOC101746277 |
| Hz_08 | HzOG207059 | 1204750 | 1211972 | 62 | -0.037 | BMORI:uncharacterized protein LOC101739635 |
| Hz_20 | HzOG210549 | 8793626 | 8804845 | 10 | 2.378 | BMORI:ankyrin-2-like |
| Hz_05 | HzOG200468 | 11331485 | 11334455 | 1100 | 2.607 | HzeaTryp125 |

| | | | | | | |
|---|---|---|---|---|---|---|
| Hz_05 | HzOG205969 | 11024388 | 11026591 | 398 | -1.525 | BMORI:protein rolling stone-like |
| Hz_09 | HzOG200434 | 5336767 | 5340510 | 49473 | 1.662 | HzeaTryp080 |
| Hz_12 | HzOG213728 | 2492271 | 2493968 | 92 | -2.655 | Lysozyme2 |
| Hz_18 | HzOG200491 | 7258652 | 7273628 | 23359 | -2.126 | HzeaChym113 |
| Hz_10 | HzOG212206 | 4686972 | 4689601 | 90 | -1.85 | BMORI:neurogenic locus notch homolog protein 1-like |
| Hz_16 | HzOG208703 | 2938423 | 2940543 | 4368 | 2.102 | BMORI:uncharacterized protein LOC101742843 |
| Hz_04 | HzOG214963 | 2912413 | 2912781 | 577 | -1.655 | DMELA:Q9W2E6 Poor Imd response upon knock-in |
| Hz_13 | HzOG207750 | 3758456 | 3759224 | 282 | -2.023 | BMORI:1-acylglycerophosphocholine O-acyltransferase 1-like |
| Hz_22 | HzOG216029 | 2026650 | 2028727 | 213 | -1.888 | BMORI:eukaryotic initiation factor 4E-1 |
| Hz_20 | HzOG207384 | 4887049 | 4889094 | 930 | -0.763 | BMORI:mRNA transport regulator 3 |
| Hz_21 | HzOG214644 | 4468459 | 4476980 | 1566 | 1.518 | BMORI:alpha-tocopherol transfer protein-like |
| Hz_28 | HzOG200286 | 3291413 | 3293452 | 136 | -0.056 | HzeaUGT33B1A |
| Hz_30 | HzOG211947 | 6284853 | 6307582 | 834 | -0.91 | BMORI:uncharacterized protein LOC101736183 |
| Hz_16 | HzOG210085 | 10926011 | 10928082 | 43 | -3.554 | BMORI:alpha-tocopherol transfer protein-like |

**Table S8.** Phenotypic results from the trypsin inhibition and leaf tissue incorporation lab assay.

| Population | Leaf Tissue | Inhibitor | n | Mean Weight (mg) | Weight sd |
|---|---|---|---|---|---|
| Susceptible | Non-expressing | None | 143 | 157.6 | 44.00768 |
| Susceptible | Non-expressing | Buffer control | 199 | 154.5 | 45.63742 |
| Susceptible | Non-expressing | 1X TLCK | 149 | 153 | 49.35163 |
| Susceptible | Non-expressing | 2X TLCK | 77 | 140.9 | 21.64248 |
| Susceptible | Non-expressing | 5X TLCK | 79 | 107.4 | 35.44576 |
| Susceptible | Non-expressing | 10X TLCK | 75 | 48.1 | 27.11194 |
| Susceptible | Cry1Ab | None | 141 | 19.7 | 13.54064 |
| Susceptible | Cry1Ab | Buffer control | 201 | 20.7 | 13.64687 |
| Susceptible | Cry1Ab | 1X TLCK | 151 | 16.7 | 12.18457 |
| Susceptible | Cry1Ab | 2X TLCK | 80 | 10.8 | 8.409723 |
| Susceptible | Cry1Ab | 5X TLCK | 80 | 5.1 | 3.643378 |
| Susceptible | Cry1Ab | 10X TLCK | 79 | 3.4 | 3.113071 |
| Resistant | Non-expressing | None | 69 | 105.6 | 50.99504 |
| Resistant | Non-expressing | Buffer control | 91 | 113.7 | 47.50252 |
| Resistant | Non-expressing | 1X TLCK | 71 | 120.5 | 35.29222 |
| Resistant | Non-expressing | 2X TLCK | 70 | 95.5 | 39.89744 |
| Resistant | Non-expressing | 5X TLCK | 72 | 57.7 | 27.7838 |
| Resistant | Non-expressing | 10X TLCK | 76 | 34.3 | 18.44673 |
| Resistant | Cry1Ab | None | 71 | 76.8 | 35.01373 |
| Resistant | Cry1Ab | Buffer control | 92 | 81.1 | 32.23311 |
| Resistant | Cry1Ab | 1X TLCK | 72 | 76.5 | 26.88142 |
| Resistant | Cry1Ab | 2X TLCK | 76 | 60.3 | 21.48516 |
| Resistant | Cry1Ab | 5X TLCK | 72 | 36.5 | 17.48786 |
| Resistant | Cry1Ab | 10X TLCK | 79 | 17.4 | 7.230397 |

**Table S9.** Pairwise post hoc comparison of treatment group effects in trypsin inhibition assays. All comparisons are between inhibitor treatments (none, buffer only, or TLCK) within a single Cry toxin treatment (Cry1Ab, or control for Cry1Ab) and population (susceptible, or resistant).

| Group A | Group B | Estimate | DF | p |
|---|---|---|---|---|
| Susceptible population - control for Cry1Ab | | | | |
| 10X | 1X | 0.0143 | 1160 | <.0001 |
| 10X | 2X | 0.0137 | 1160 | <.0001 |
| 10X | 5X | 0.0115 | 1160 | <.0001 |
| 10X | CL | 0.0143 | 1160 | <.0001 |
| 1X | 2X | -0.0006 | 1160 | 1 |
| 1X | 5X | -0.0028 | 1160 | 0.0005 |
| 1X | CL | 0.0001 | 1160 | 1 |
| 2X | 5X | -0.0022 | 1160 | 0.0391 |
| 2X | CL | 0.0006 | 1160 | 1 |
| 5X | CL | 0.0028 | 1160 | 0.0002 |
| Susceptible population - Cry1Ab | | | | |
| 10X | 1X | 0.236 | 1160 | <.0001 |
| 10X | 2X | 0.203 | 1160 | <.0001 |
| 10X | 5X | 0.0987 | 1160 | 0.0002 |
| 10X | CL | 0.247 | 1160 | <.0001 |
| 1X | 2X | -0.0325 | 1160 | <.0001 |
| 1X | 5X | -0.137 | 1160 | <.0001 |
| 1X | CL | 0.0116 | 1160 | 0.0075 |
| 2X | 5X | -0.104 | 1160 | <.0001 |
| 2X | CL | 0.044 | 1160 | <.0001 |
| 5X | CL | 0.148 | 1160 | <.0001 |
| Resistant population - Control for Cry1Ab | | | | |
| 10X | 1X | 0.0208 | 761 | <.0001 |
| 10X | 2X | 0.0187 | 761 | <.0001 |
| 10X | 5X | 0.0118 | 761 | <.0001 |
| 10X | CL | 0.0203 | 761 | <.0001 |

| | | | | |
|---|---|---|---|---|
| 1X | 2X | -0.0022 | 761 | 0.0122 |
| 1X | 5X | -0.009 | 761 | <.0001 |
| 1X | CL | -0.0005 | 761 | 1 |
| 2X | 5X | -0.0069 | 761 | <.0001 |
| 2X | CL | 0.0017 | 761 | 0.1073 |
| 5X | CL | 0.0085 | 761 | <.0001 |
| Resistant population - Cry1Ab | | | | |
| 10X | 1X | 0.0443 | 761 | <.0001 |
| 10X | 2X | 0.0408 | 761 | <.0001 |
| 10X | 5X | 0.03 | 761 | <.0001 |
| 10X | CL | 0.0451 | 761 | <.0001 |
| 1X | 2X | -0.0035 | 761 | 0.0068 |
| 1X | 5X | -0.0143 | 761 | <.0001 |
| 1X | CL | 0.0007 | 761 | 1 |
| 2X | 5X | -0.0108 | 761 | <.0001 |
| 2X | CL | 0.0043 | 761 | 0.0001 |
| 5X | CL | 0.015 | 761 | <.0001 |

**Table S10**. Primers for ddPCR copy number variant analysis of the Trypsin 77 target gene and ATP dependent DNA helicase control gene.

| Assay | Primer pair | Sequence |
|---|---|---|
| Control gene | Forward | CATGTTCCCAAATGTGCCTATC |
| | Reverse | AACAAGGCAACCAGGAATATTAAG |
| Target gene (Trypsin 77) | Forward | TGAGGGTGGATTGTGAAGTTAG |
| | Reverse | GCGTCAGAACACCGTCTATT |

**Supplementary Figures**

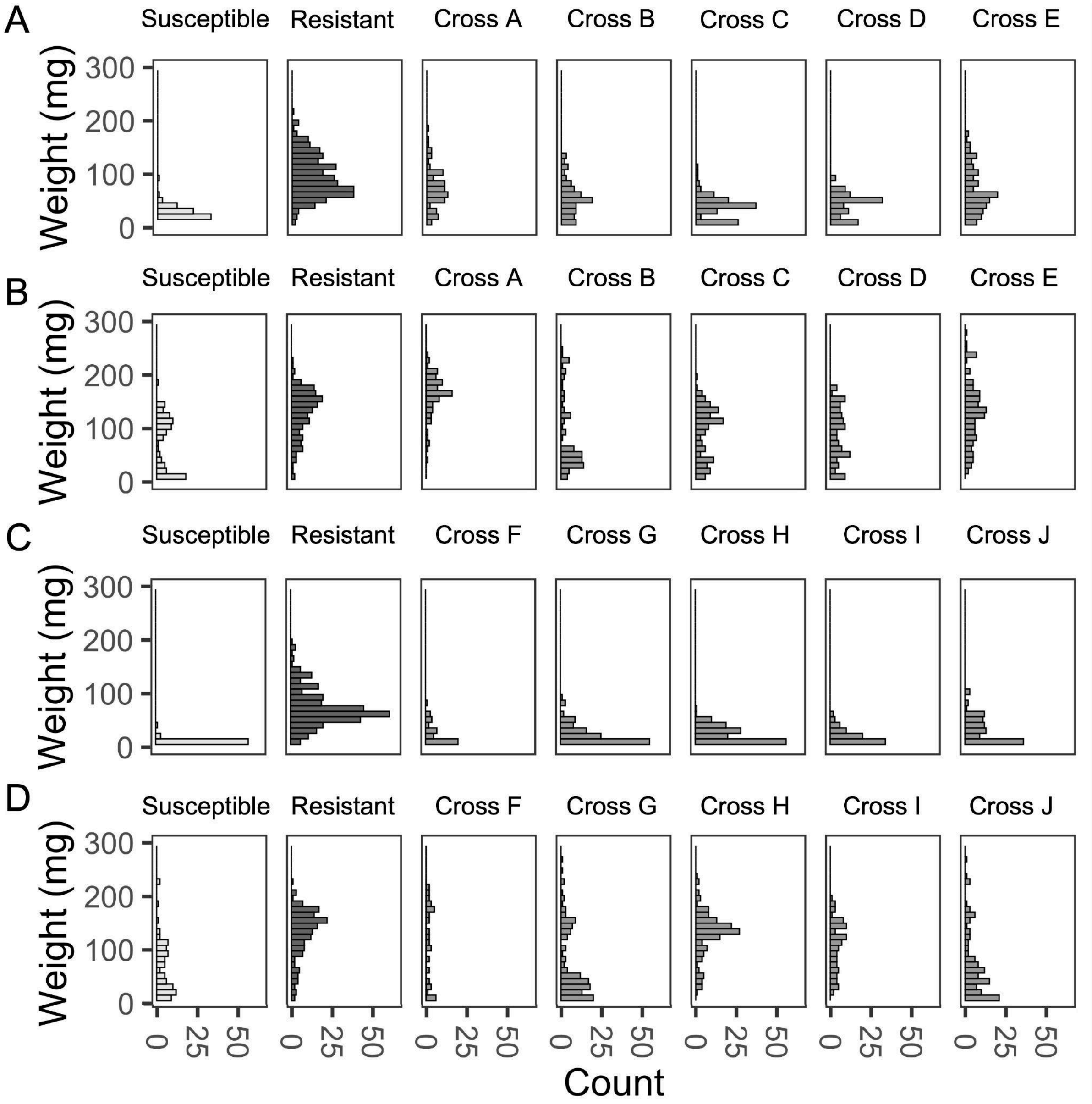

**Figure S1**. Weight histograms after seven days in a laboratory leaf tissue incorporation assays for Cry toxin containing and control treatments. The susceptible laboratory colony is shown in light gray, the field derived resistant *H. zea* are shown in dark gray, and the $F_2$ intercross progeny between resistant and susceptible populations are shown in medium gray and split by cross family. Weight is shown for assays with Cry1Ab expressing leaf tissue (**A**), its non-expressing control near isoline **(B)**, Cry1A.105 and Cry2Ab2 expressing leaf tissue **(C)**, and its non-expressing control near isoline **(D).**

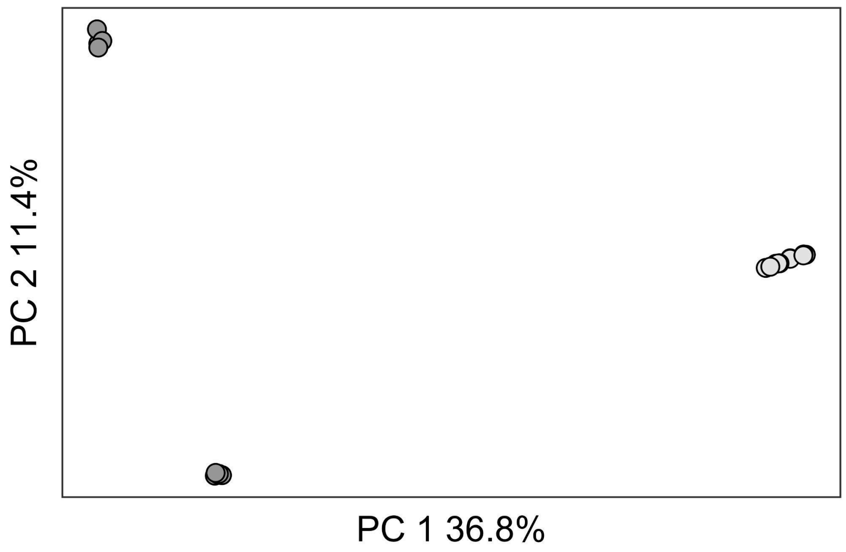

**Figure S2.** Principal component analysis for resistant and susceptible cross parents using genome wide SNPs with Plink. Resistant (dark gray and) susceptible (light gray) individuals are separated by PC 1 which explains 36.8% of variance while PC 2 captures differences between field samples and explains 11.4% of variance.

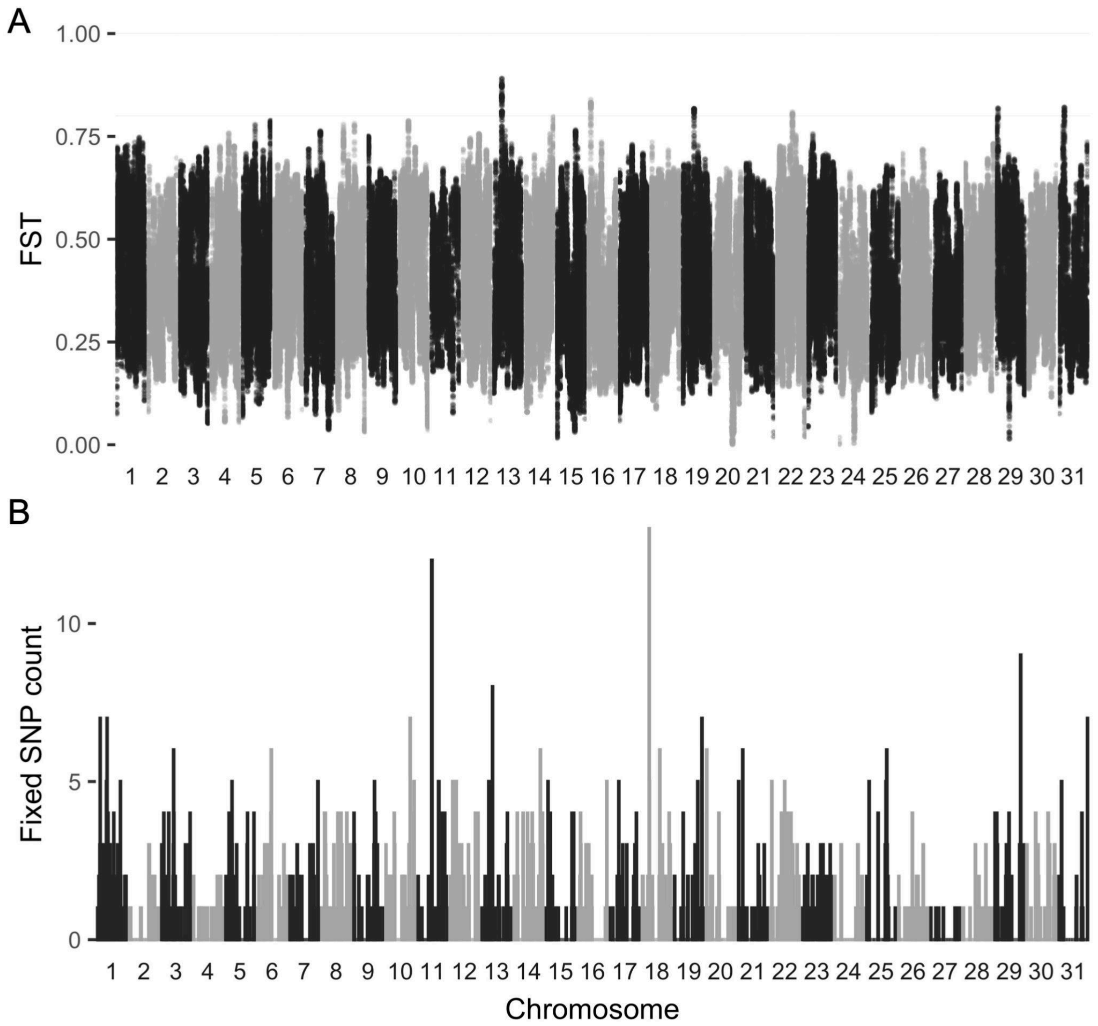

**Figure S3.** Genome wide differentiation between resistant field and susceptible lab colony individuals. **A.** Divergence measured with Weir and Cockerham's windowed $F_{ST}$ across 40kb windows with a 10kb step plotted across all 31 chromosomes. Divergence between these populations is high (mean window $F_{ST}$ = 0.405) and variance is wide (window $F_{ST}$ range = -0.045 - 0.891). No windows fell above the genome wide $zF_{ST}$ significance threshold of 6. **B.** Density of variant SNPs fixed between resistant and susceptible individuals in 100kb windows across the 31 chromosomes.

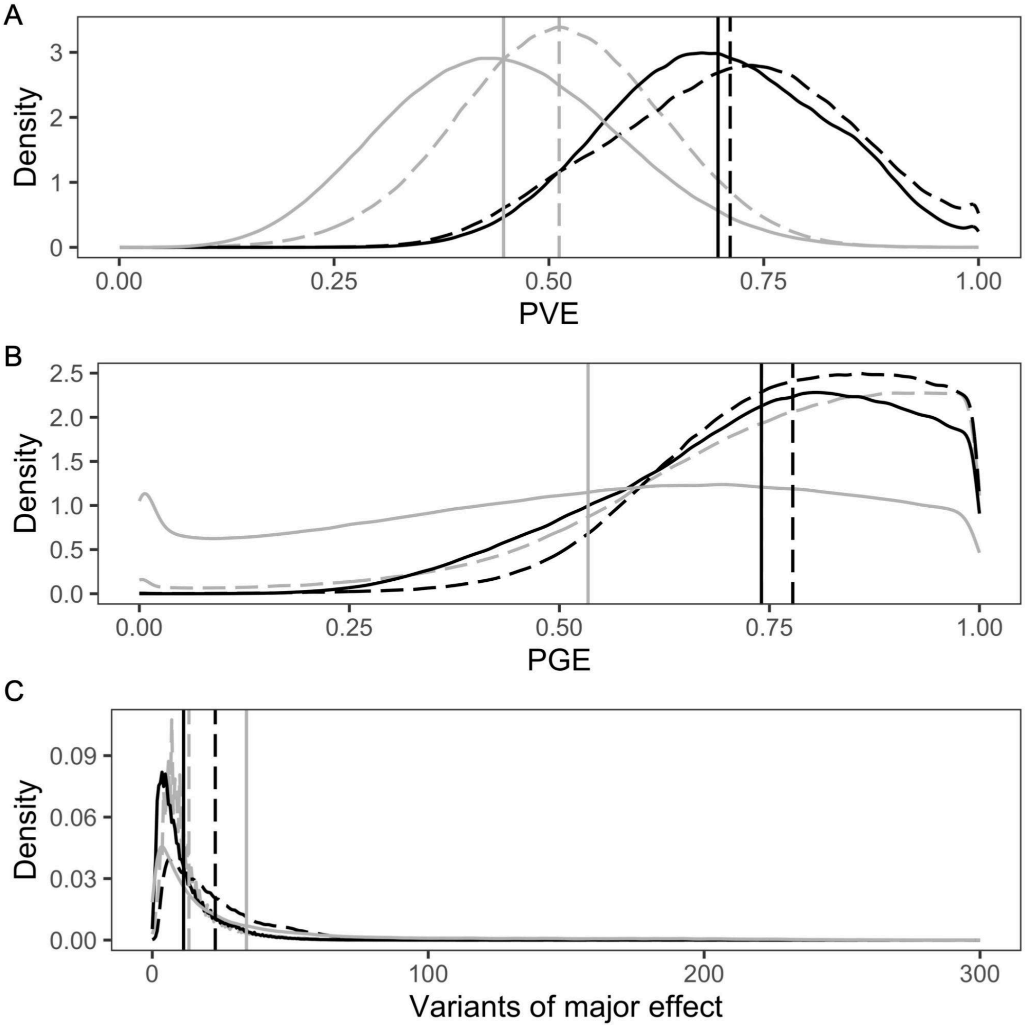

**Figure S4**. BSLMM hyperparameter estimate distributions for proportion of phenotypic variance explained by the whole panel of genomic variants PVE **(A)**, the proportion of that genetic variance explained by loci of major effect PGE **(B)**, and the number of variants of major effect **(C)**. The estimates for the two Cry resistance traits are shown in dark gray while the estimates for the two control growth traits are shown in light gray. The Cry1Ab resistance trait and its associated control is shown with a solid line while the Cry1A.105+Cry2Ab2 trait and its associate control are shown with dashed lines.

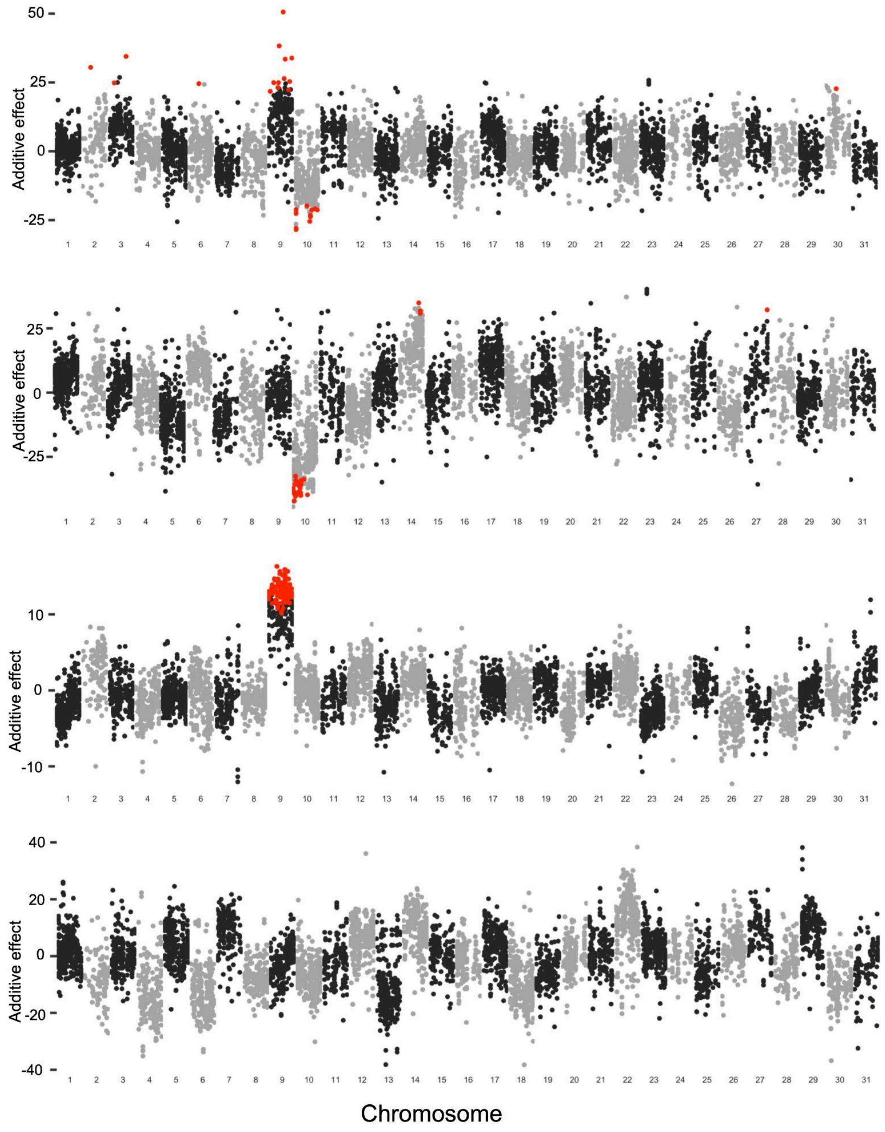

**Figure S5**. Unsmoothed association of genome wide markers and weight gain phenotype after seven days of feeding on Cry toxin containing and control diets. The additive effect (beta from LMM) is the weight gain in mg associated with the presence of a single resistant population allele for markers on 31 *H. zea* chromosomes. The genotype phenotype association is shown for **A.** weight gain after seven days of exposure to Cry1Ab, **B.** weight gain on the non-toxin near isoline control diet for Cry1Ab, **C.** weight gain after seven days of exposure to Cry1A.105 + Cry2Ab2, **D.** weight gain on the non-toxin near isoline control diet for Cry1A.105 + Cry2Ab2.

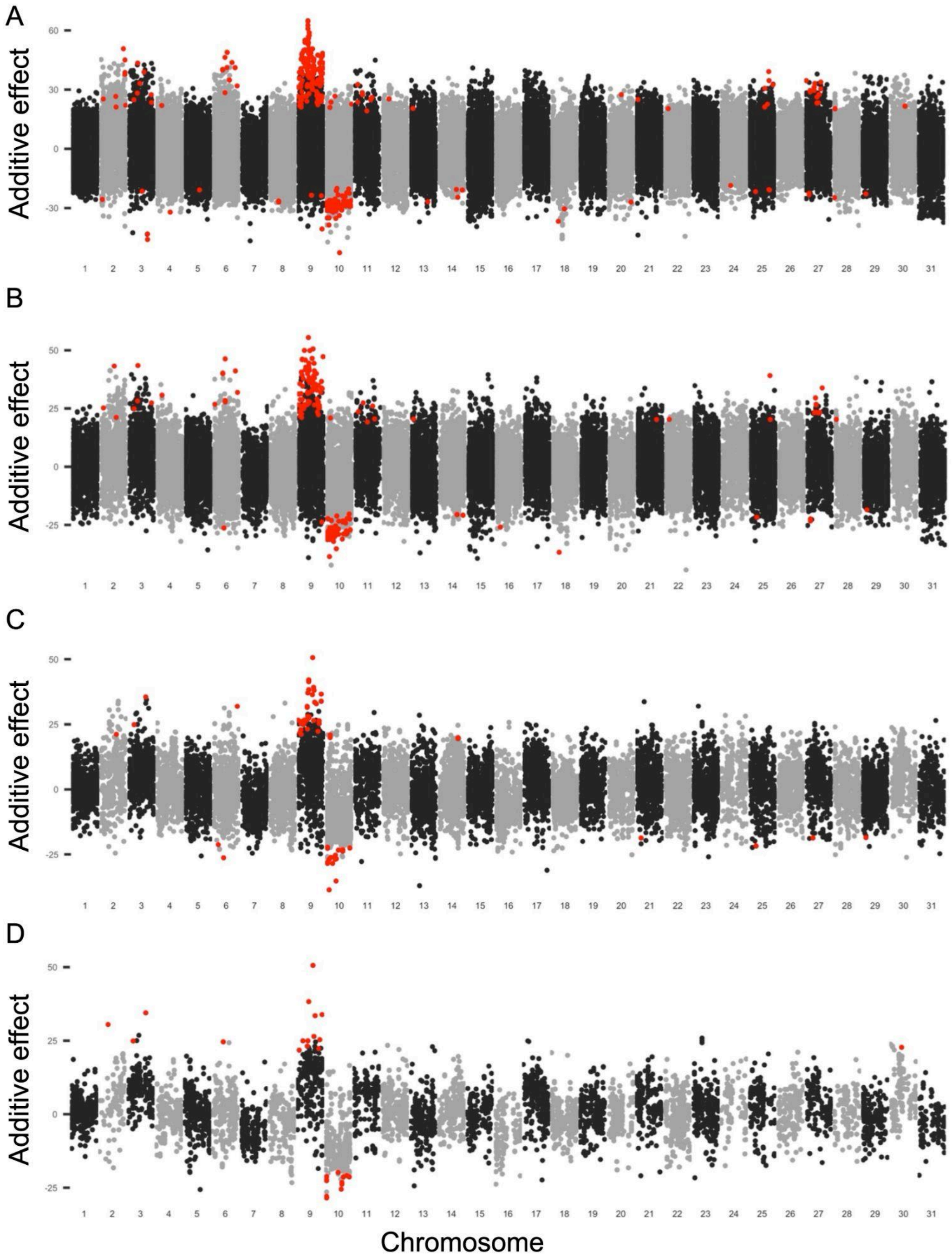

**Figure S6**. Genotype phenotype effect direction determination with different allele count filtering thresholds between resistant and susceptible populations. **A.** Shows the full dataset, while data subsets with increasingly stringent allele count filtering thresholds are shown below. The thresholds used on the lower panels are **B.** > 5, **C.** >10, and **D.** >15.
